# Enhancer and super-enhancer landscape in polycystic kidney disease

**DOI:** 10.1101/2021.11.19.469306

**Authors:** Ronak Lakhia, Abheepsa Mishra, Laurence Biggers, Venkat Malladi, Patricia Cobo-Stark, Sachin Hajarnis, Vishal Patel

## Abstract

Widespread aberrant gene expression is pathological hallmark of polycystic kidney disease (PKD). Numerous pathogenic signaling cascades, including c-Myc, Fos, and Jun are transactivated. However, the underlying epigenetic regulators are poorly defined. Here we show that H3K27ac, a histone modification that marks active enhancers, is elevated in mouse and human ADPKD samples. Using comparative H3K27ac ChIP-Seq analysis, we mapped >16000 active intronic and intergenic enhancer elements in *Pkd1*-mutant mouse kidneys. We find that the cystic kidney epigenetic landscape resembles that of a developing kidney, and >90% of upregulated genes in *Pkd1*-mutant kidneys are co-housed with activated enhancers in the same topologically associated domains. Furthermore, we identify an evolutionarily-conserved enhancer cluster downstream of the *c-Myc* gene and super-enhancers flanking both *Jun* and *Fos* loci in mouse and human ADPKD models. Deleting these regulatory elements reduces *c-Myc*, *Jun*, or *Fos* abundance and suppresses proliferation and 3D cyst growth of *Pkd1*-mutant cells. Finally, inhibiting glycolysis and glutaminolysis or activating *Ppara* in *Pkd1*-mutant cells lowers global H3K27ac levels and on *c-Myc* enhancers. Thus, our work suggests that epigenetic rewiring mediates the transcriptomic dysregulation in PKD, and the regulatory elements can be targeted to slow cyst growth.

## INTRODUCTION

Autosomal Dominant Polycystic Kidney Disease (ADPKD), primarily caused by mutations in the *PKD1* or *PKD2* genes, affects nearly 12 million individuals worldwide and equally affects individuals irrespective of gender or race^1^. Fifty percent of individuals with ADPKD develop kidney failure requiring dialysis or kidney transplant ^2^. The clinical hallmark of ADPKD is the massive bilateral kidney enlargement due to numerous kidney tubule-derived cysts. These cysts are lined by rapidly proliferating and metabolically deranged tubular epithelial cells^3–7^, fueling their progressive and relentless growth. Additionally, there is substantial interstitial inflammation and fibrosis, which also contribute to the decline in kidney function^8^. Tolvaptan, a vasopressin receptor 2 antagonist, slows the rate of kidney function decline and is the only FDA-approved treatment ^9, 10^. There are other emerging therapeutic modalities^11–14^, but ADPKD pathogenesis is still incompletely understood, and there is a dire need for uncovering new drug targets.

Enhancers are dynamic cis-regulatory DNA elements (CREs), approximately 200–2,000 bp in length, that help shape cellular and tissue identity by fine-tuning transcriptomic makeup^15^. Active enhancers are marked by the transcriptionally-permissive H3K27 acetylation (H3K27ac) histone modification and accessible chromatin that allows the binding of sequence and cell type-specific transcription factors, transcriptional co-activators, and RNA polymerase II (RNAP II). A subset of CREs is referred to as super-enhancers^16–18^. These span large genomic regions (several kb in length) and exhibit unusually high transcriptional factor binding density and H3K27ac histone modifications. Active enhancers/super-enhancers physically interact with gene promoters, often over long genomic distances, to regulate gene transcription^15^. Unlike promoters, enhancers/super-enhancers can be present upstream or downstream of their target genes and are capable of activating gene transcription in any orientation. The enhancer/super-enhancer landscape is tissue-specific, contributing to each organ’s unique transcriptomic output. Enhancers are also dynamically regulated, facilitating gene expression switches accompanying tissue developmental stage. With regards to diseases, enhancer biology is best studied in cancer, where many tumor-specific enhancers and super-enhancers have been reported. Moreover, targeting enhancers has shown early promise as a novel method to reduce pathogenic gene expression. However, barring a few exceptions^19, 20^, enhancer impact on most kidney diseases still remains poorly defined.

Large-scale transcriptomic dysregulation is a pathological hallmark of ADPKD. Numerous pro-proliferative and pathogenic signaling cascades, including c-Myc, Fos, Jun, and cAMP ^21–23^, are transactivated in cystic kidneys. Thus, the goal of this study was to define the epigenetic mechanisms that underlie this widespread aberrant gene expression. In this work, we perform a comparative H3K27ac ChIP-Seq analysis and provide a detailed enhancer/super-enhancer map of cystic kidneys. We find that mice and human ADPKD kidneys bear a similar enhancer signature. Moreover, we dissect and characterize a series of enhancers flanking the pathogenic c-*Myc*, *Jun,* and *Fos* genes. Finally, our studies suggest that enhancer rewiring is partly fueled by the metabolic reprogramming observed in ADPKD kidneys. Together, the epigenetic map unveiled by our studies has uncovered a series of potential new targets to slow cyst growth.

## METHODS

### Mice

Ksp^Cre^, *Pkd1*^F/F^, *Pkd1*^RC/RC^, Pkhd1^Cre^ and *Pkd2*^F/F^ mouse strains have been previously described^24–28^. The Ksp^Cre^; *Pkd1*^RC/RC^ mice were bred with *Pkd1*^F/F^ mice to obtain Cre-negative*; Pkd1*^F/RC^ (control) and Ksp^Cre^*; Pkd1*^F/RC^ (*Pkd1-*mutant) mice. Kidneys were harvested at the ages noted in results. All mice were maintained in C57Bl/6 background. For ChIP experiments, each biological replicate consisted of four pooled kidneys from two mice of the same sex. Equal males and females were used in all studies. All experiments were performed according to the guidelines and approval of the UT Southwestern Institutional Animal Care and Use Committee.

### Human Specimens

Human kidney specimens (ADPKD and NHK) were provided by the PKD Biomarkers and Biomaterials Core at the Kansas PKD center at the Kansas University Medical Center (KUMC). Kidneys were procured through the Bio-specimen Resource Core at the University of Kansas Cancer Center. Harvested kidneys were instantly sealed in sterile bags in the operating room, submerged in ice, and sent to the laboratory for processing. All surgical and processing protocols followed federal regulations and were approved by the Institutional Review Board at KUMC.

### Cell culture

*Pkd1*^RC/+^ and *Pkd1*^RC/-^ cells were generated and maintained in previously described epithelial media with 2% fetal bovine serum (FBS) at 37°C^29^. Briefly, two kidneys were harvested from a 14-day old *Pkd1*^RC/F^ male mouse and minced to 1mm cubes. The tissue was incubated for 40 min in DMEM containing 5% Collagenase (Sigma #C1639, USA) at 37°C with intermittent agitation to create a single-cell suspension. Cells were strained using a 40-micron cell strainer and incubated with Biotinylated Dolichos Biflorus Agglutinin (DBA) (Vector labs #B-1035) for 1 hour. DBA positive cells were isolated using a CELLection Biotin binder kit (Invitrogen #11533D) and placed in culture media overnight. Next, cells were immortalized using the SV40 T Antigen Cell Immortalization Kit (Alstem #CILV01) and underwent a clonal expansion in 96 well plates. Clones were screened for SV40 marker by genotyping for SV40, and one clone was selected for infection with an adenovirus that expresses Cre recombinase (Vector Biolabs #1779). Infected cells subsequently underwent clonal expansion. Genotyping for the recombined *Pkd1* allele confirmed *Pkd1*^RC/-^ genotype.

### Chromatin Immunoprecipitation (ChIP)

ChIP was performed using Simple ChIP Enzymatic Chromatin IP Kit (Cell Signaling #9005). Briefly, tissues were minced and then cross-linked with 37% formaldehyde for 20 minutes. Specimens were then neutralized with glycine, washed with PBS and protease inhibitor cocktail, and then underwent lysis. Fragmentation was performed by micrococcal nuclease, and the lysate was obtained via mild sonication. Clarified lysate was incubated with 1 mg of H3K27ac antibody (Abcam #ab4729) or IgG overnight on a rotating platform kept at 4°C. Before adding the antibody, 20 ul of the lysate (2%) was removed (input). To the enriched lysate, 30 ul of ChIP-grade Protein G was added and put on a rotating platform for 3 h at 4°C. The chromatin-complex bound to beads was washed three times with low salt wash buffer and once with high salt wash buffer. The chromatin complex was then eluted from beads at 65°C with gentle shaking for 30 minutes. The protein-DNA cross-links were reversed, and the DNA was purified using DNA purification spin columns and recovered efficiently. The eluted DNA was quantified using Qubit™ dsDNA HS Assay Kit (Invitrogen #Q32851, USA). The H3K27ac enriched regions were analyzed by quantitative real-time PCR and ChIP-seq. (**Suppl. Table 2 and 3**). KAPA HTP Library Preparation Kit was used to prepare libraries and samples were run on Illumina HiSeq 2500.

### Analysis of ChIP-seq Data

The raw reads were aligned to the mouse reference genome (GRCh38/mm10) using default parameters in BWA v0.7.12^30^. The aligned reads were subsequently filtered for quality, and uniquely mappable reads were retained for further analysis using Samtools version 1.6 and Sambamba version 0.6.6 ^31, 32^. Library complexity was measured using BEDTools version 2.26.0 and meets ENCODE data quality standards^33, 34^. Relaxed peaks were called using MACS version 2.1.0 with a p-value of 1×10^-2^ for each replicate, pooled replicates’ reads, and pseudoreplicates^35^. Peak calls from the pooled replicates that were either observed in all replicates or both pseudoreplicates were used for subsequent analysis.

### Identification of variant enhancer loci

To identify enhancers, we first mapped transcription start sites (TSS) for protein-coding genes from Gencode version 15 annotations using MakeGencodeTSS (https://github.com/sdjebali/MakeGencodeTSS)36. Potential enhancers were defined as peaks that were > 2kb away from known TSS, protein-coding genes. To detect differentially bound enhancers, we used the R package DiffBind version 2.2.12 with a p-value < 0.05 and default parameters.

### Identification of super-enhancers loci

Super-enhancers were called using the ROSE version 0.1 package using the universe of H3K27ac peaks called for each mouse model^17, 37^. Differential super-enhancers were assessed using a universe of super-enhancers created by using the R package DiffBind version 2.2.12. Quantification of differential super-enhancers was assessed from using the master list and counting the overlapping differential enhancers previously identified using BEDTools version 2.26.0^33^. We identified a differential super-enhancer if a given mouse model had at least one enhancer present. The gained and lost super-enhancers were identified to have more differential enhancers.

### Generating Heatmaps and Metagenes

Enrichment of ChIP-seq signal was assessed by generating heatmaps and metagenes. For heatmaps and metagenes of ChIP-seq intensities, we used deepTools^38^. Read abundances were generated using ’computeMatrix’ around the peak center and subsequently used to generate heatmaps and metagenes using ’plotHeatmap’ and ’plotProfile’, respectively.

### Peak Annotations and Motif Enrichment

Peak annotations were identified by using the R 3.3.4 package ChIPseeker version1.1.18 using ’TxDb.Mmusculus.UCSC.mm10.knownGene’ and ’org.Mm.eg.db’ and default parameters^39^. To identify enriched motifs, we performed motif enrichment analysis by using HOMER version 4.9^40^.

### Analysis of Embryonic ChIP-seq data

Embryonic data sets with alignments and peak files for H3K27ac ChIP-seq experiments (ENCSR057SHA; E15.5) and (ENCSR178MLS; P0) were extracted from ENCODE. The datasets were analyzed following the same method as our H3K27ac ChIP-seq datasets.

### SnATAC-seq Analysis

Publicly-available dataset GSE157079 was downloaded from Gene expression omnibus^41^. Minimum peak accessibility of each cell type was determined by comparing peak counts per cell type to a null peak count of the same total size using a one-tailed Fisher exact test. Those cell types with a p-value <0.05 for a peak were considered accessible at that peak. Differential peak accessibility of cell types was scored by comparing the peak counts per cell type to those of all other cell types using a two-tailed Fisher exact test with Benjamini-Hochberg p-value correction. If a cell-type pair comparison for a peak has an adjusted p-value <0.05, then the upregulated cell type’s score was incremented.

### TAD-data association

RNA-seq and ChIP-seq datasets were binned into the topologically associated domains (TAD) of mouse embryonic stem cells (mESC)^42^. The midpoint of each transcript or peak was used to determine the TAD assignment.

### RNA extraction and qPCR

RNA extraction was performed with miRNeasy Kit (Qiagen #217004). Superscript III (Invitrogen) was used for the synthesis of first-strand cDNA from mRNA. qPCR was performed using the iQ SYBR Green Supermix (Bio-Rad) using the CFX ConnectTM Real-time PCR detection system. All samples were analyzed with technical replicates. Ribosomal 18S was used to normalize mRNA expression.

### Generation of CRISPR-KO cell lines

SgRNAs were designed to flank enhancers and cloned into pX330mCherry and SpCas9-2A-GFP as previously described **(Suppl. Table 4)**^43, 44^. Cells were transfected using Lipofectamine 3000 Reagent (Invitrogen) and underwent FACS sorting 72 hours after transfection to select for double-positive cells and subsequent single-cell plating in 96 well plates. Clonal cell lines were genotyped (**Suppl Table 5**) to assess for deletion of each respective enhancer. In each experiment, clonal cell lines which underwent transfection with sgRNAs but did not develop deletion of enhancers were utilized as control cell lines for subsequent experiments.

### Western blot

Total protein was extracted with Tissue Protein extraction reagent (T-PER, Thermofisher Scientific, USA) + Protease inhibitor cocktail (Pierce #A32961). Protein concentration was measured using Bradford’s reagent. Ten to twenty micrograms of denatured protein was loaded onto a 4-20% TGX stain-free polyacrylamide gel (Bio-Rad, USA) or NuPAGE 3-8% Tris acetate gel and subsequently transferred onto pre-cut nitrocellulose membranes. The membrane was blocked in 5% milk for 1h followed by incubation with primary antibodies: c-Myc (Abcam ab185656; 1:1000), Fos (Cell Signaling 2250S; 1:1000), Jun (Cell Signaling 9165; 1:1000), CREB (Cell Signaling 9197S; 1:1000), STAT1 (Cell Signaling 14994S; 1:1000), PC1(7e12) (Santa Cruz #SC-130554 1:1000), PC2 (Baltimore PKD Center) or Actin (Sigma a3854;1:40,000) overnight or 30 min at 4°C. The blots were washed with TBST and probed with Goat anti-Rabbit-HRP conjugated secondary antibody for 1h. The blots were washed three times with TBST prior to development with SuperSignal West Femto, SuperSignal West Dura Extended Dilution Substrate, or ECL.

### Alamar Blue assay

1×10^4^ cells were plated in 24-well plates. The following day, fresh media containing 10% Alamar Blue reagent was added. Absorbance at 570 and 600 nm was measured using BioTek Synergy H1 microplate reader 12 hours post-incubation, and percentage reduction was determined.

### 3D Cystogenesis Assay

Matrigel basement membrane matrix (Corning, #354234) was thawed on ice for 30 min. Micropipette tips and 8-well chamber slides were pre-cooled at 4 degrees. 8- well LAB-TEK II chamber slides w/ cover glass (Corning # 155409) were placed on ice and coated with 25 μl 100% matrigel and then placed at 37 degrees. Cells were trypsinized, counted, and diluted to 1.5×10^4^ cells/ml in epithelial media. Matrigel was diluted in epithelial media to achieve a final concentration of 4%. The cell suspension and 4% matrigel were mixed (1:1) to achieve a final concentration of 2% matrigel. 300 μl of cell/matrigel mix was added to each well. 7 days post-seeding, 20x images of cysts were obtained using light microscopy and quantified with Image J software. For each experiment, 100 cysts were quantified. Each experiment was repeated 3 times (biological replicates).

### Histone extraction

Histones were extracted via an acid-extraction method. Briefly, samples were weighed, cut, and minced with a blade. The minced tissue was then transferred to a Dounce homogenizer and re-suspended in TEB buffer (PBS with 0.5% Triton X 100, 2 mM PMSF, and 0.02% NaN3) at 0.5 ml per 100 mg tissue and disaggregated by 50 strokes. The homogenate was transferred to a 2 ml Eppendorf tube and centrifuged at 10,000 rpm for 1 min at 4°C. The cells were harvested and centrifuged at 1000 rpm for 5 minutes at 4°C. The cell pellet was re-suspended in TEB buffer at 10^7^ cells/0.5ml and lysed on ice for 10 minutes via vortex. The cell lysates were centrifuged at 10,000 rpm for 1 minute at 4°C. The supernatant was removed, and the cell/tissue pellet was re-suspended in 3 volumes (approx. 100 µl/10^7^ cells or 100 mg tissues) of extraction buffer (0.5N HCl + 10% glycerol) and was incubated on ice for 30 min. The cell lysate was centrifuged at 12,000 rpm for 5 minutes at 4°C, and the supernatant was transferred to a new vial. To the supernatant, approximately eight volumes (0.3 ml/10^7^ cells or 100 mg tissues) of acetone was added and stored at –20°C overnight. The supernatant-acetone mixture was centrifuged at 12,000 rpm for 5 minutes and air-dried for 10-20 min. The pellet was dissolved in distilled water (30 µl/10^7^ cells or 100 mg tissues). Bradford assay was used to quantify histone abundance for subsequent use in ELISA assay.

### Histone H3 acetyl Lys27 ELISA

The Histone H3 acetyl Lys27 ELISA (H3K27ac) kit (Active Motif, #53116) was used to detect H3K27ac levels from crude acid extracted histone preparations. Briefly, the sandwiched ELISA kit uses a 96-well strip coated with H3K27ac monoclonal antibody for capturing histone (H3) from samples followed by incubation with anti-mouse H3K27ac primary antibody (1:500). A secondary antibody conjugated with HRP (1:500) and developing substrate is used for colorimetric detection at 450 nm. A recombinant H3K27ac protein provided in the kit was used to plot a reference standard curve to quantify the amount of H3K27ac present in each sample.

### Immunostaining

Paraffin-embedded sections were used for mouse kidney tissues. In the case of human kidney tissues, frozen sections were used. Both anti-H3K27ac (Abcam #ab4729) and Alexa Fluor anti-Rabbit 594 secondary antibodies were used at 1:500 dilution for immunostaining experiments.

### Metabolism Experiments

*Pkd1*^RC/-^ cells were grown to 60% confluency and then incubated with 20 μM 2-DG for 24 hours, 40 μM BPTES for 48 hours, or 40 μM of WY-14643 for 72 hours. Corresponding vehicle controls also underwent equivalent incubation times before harvest for ChIP, ELISA, and Western blot analysis.

### Antisense Oligonucleotide Treatment

Antisense oligonucleotides (ASOs) were designed and obtained from Qiagen. 2×10^^5^ cells were seeded in a 6 well plate, and the following day media was changed. Lipofectamine 3000 was used to transfect 40 nM scramble ASO or *Junos* ASO (cocktail of 3 ASOs targeting 3 different regions of the *Junos* transcript at final total concentration of 40 nM). Fluorescent microscopy was performed after 24 hours to confirm adequate delivery of ASO by visualization of FAM. Cells were harvested after 72 hours for subsequent molecular analysis.

### Data Availability

The ChIP-seq data are available at Geo Expression Omnibus GSE189153.

### Statistical Analysis

For statistical analysis, a two-tailed Student’s t-test for pairwise comparison was performed. For 3D cyst assay and Alamar blue experiments, a nested t-test was performed. *P*<0.05 was considered statistically significant.

## RESULTS

### H3K27ac histone modification is increased in mouse and human ADPKD samples

Active enhancers are marked by H3K27ac, an acetylation of lysine residue (27^th^ position) of the DNA packing protein Histone H3^45^. Therefore, to assess the epigenetic status of cystic kidneys, we began by measuring global H3K27ac abundance in kidneys of orthologous ADPKD mouse models. We extracted histones from whole kidneys of 18-day-old Ksp^Cre^;*Pkd1*^F/RC^ (*Pkd1*-mutant), 28-day-old Pkhd1^cre^; *Pkd2*^F/F^ (*Pkd2*-KO), and their respective age-matched littermate control mice. We then used ELISA to quantify H3K27ac levels within the extracted histones. We found that H3K27ac levels were 55% higher in *Pkd1*-mutant and 17% higher in *Pkd2*-KO kidneys compared to their respective age-matched control kidneys (**Fig. 1A-B**). Moreover, staining with an anti-H3K27ac antibody also revealed a higher H3K27ac immunofluorescence signal. Quantification of random 20x images demonstrated that the H3K27ac signal was higher by 55% in *Pkd1*-mutant kidneys (**Fig. 1D-E and L**) and by 64% in *Pkd2*-KO kidneys compared to their respective control kidneys (**Fig. 1F-G and M**). We also measured H3K27ac abundance in the long-lived, slowly progressive *Pkd1*^RC/RC^ ADPKD mouse model. Compared to age-matched controls, the H3K27ac level was increased in 160-day-old *Pkd1*^RC/RC^ mice kidneys by 55 percent (**Fig. 1H, I, and N**). Notably, the predominant increase in the H3K27ac signal in each mouse model originates from the cyst epithelial cells.

**Figure 1.**
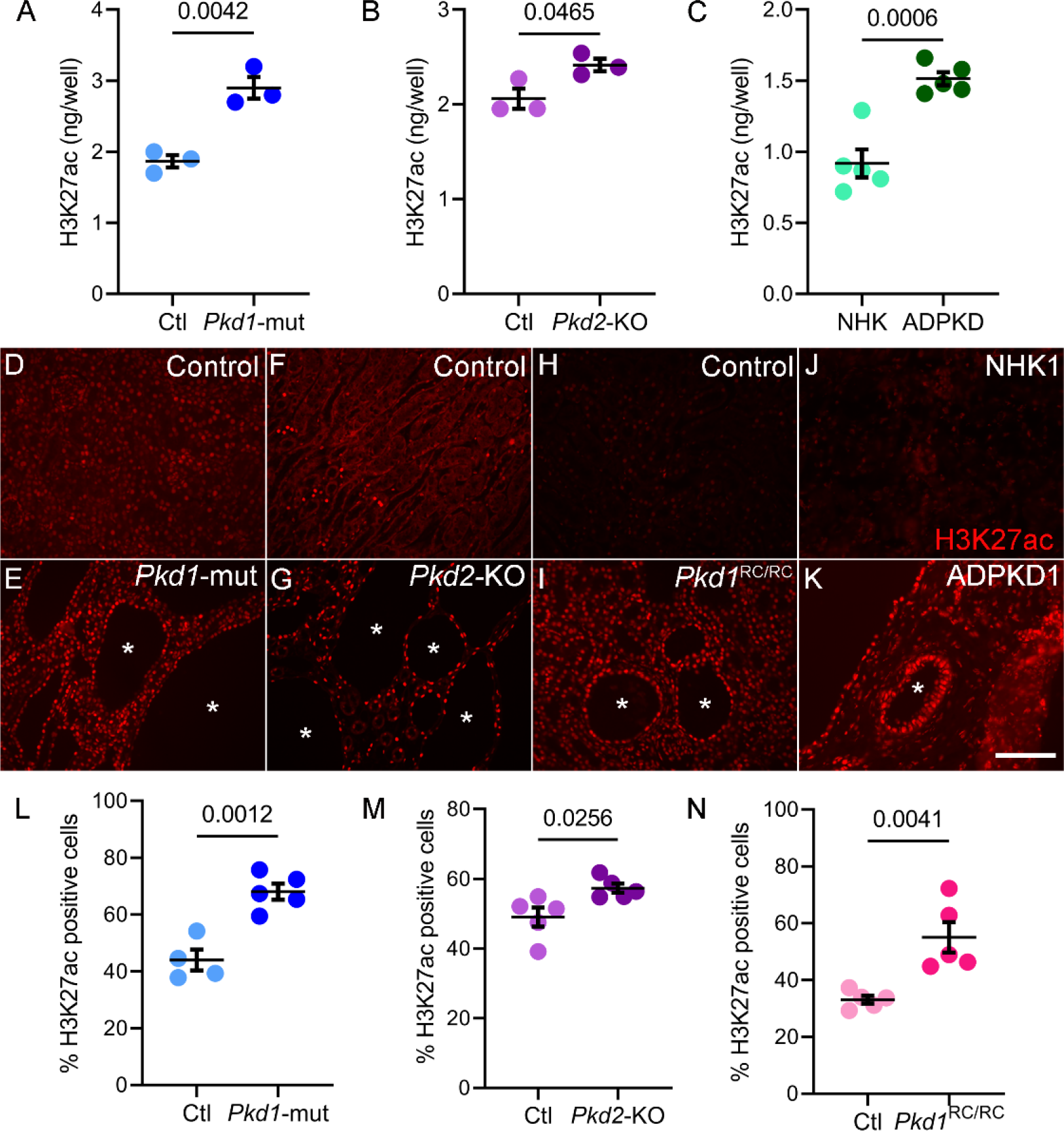
Global H3K27ac level is increased in ADPKD models. **A and B.** ELISA showing higher H3K27ac levels in 18-day-old *Pkd1*-mutant and 21-day-old *Pkd2*-KO kidneys compared to their respective age-matched control kidneys (N=3). **C.** ELISA showing higher H3K27ac levels in human ADPKD samples compared to normal human kidney (NHK) samples (N=5). **D-K** Representative images of immunofluorescence staining with anti-H3K27ac antibody in kidney sections of 18-day-old *Pkd1*-mutant, 21-day-old *Pkd2*-KO, 160-day-old *Pkd1^RC/RC^,* and human ADPKD samples compared their respective controls. **L-N.** Quantification of the H3K27ac signal using the Image J software is shown. Error bars indicate SEM; * cyst; scale bar = 100 μM; Statistical analysis: Student’s *t*-test.

To determine whether our observations in mouse models are relevant to human ADPKD, we measured H3K27ac levels using ELISA (N=5) and immunofluorescence (N=3) in normal human kidney (NHK) samples and cystic kidney samples from individuals with ADPKD. Similar to murine ADPKD models, ELISA-measured H3K27ac level was increased in human ADPKD samples by 64% compared to NHK controls (**Fig. 1C**). Moreover, immunofluorescence staining of kidney sections from a different group of NHK and ADPKD kidney samples demonstrated higher H3K27ac signal primarily enriched in cyst epithelial cells (**Fig. 1J and K and Suppl. Fig. 1**). Thus, global H3K27ac levels are increased in both mouse and human ADPKD.

### Comparative H3K27ac ChIP-Seq uncovers the enhancer landscape of cystic kidneys

Our observation of higher H3K27ac levels in ADPKD models points to substantial epigenetic rewiring in cystic kidneys. As the first step towards identifying the enhancer landscape of cystic kidneys, we immunoprecipitated and sequenced DNA (ChIP-Seq) bound to H3K27ac-modified histones in 16-day-old control and *Pkd1*-mutant kidneys (N=3, each replicate consisted of chromatin pooled from four kidneys) (**Fig. 2A**). We used 16-day-old *Pkd1*-mutant kidneys because they are mildly cystic, allowing us to identify enhancers in the early stages of cystogenesis. Principal component analysis revealed that control and *Pkd1*-mutant ChIP-Seq samples clustered separately (**Suppl Fig. 2A**). We cross-compared the H3K27ac-modified epigenome of control and cystic kidneys. In concordance with our findings of higher total H3K27ac levels, we found that 16560 regulatory elements were gained (increased H3K27ac), and only 1552 were lost (decreased H3K27ac) in *Pkd1*-mutant compared to control kidneys (**Fig. 2B**). Moreover, annotation of these regulatory elements revealed that 93% of the H3K27ac ChIP-seq peaks were located in intron or intergenic regions (**Fig. 2C**). Only 2.8% of peaks were found in promoter regions indicating that the gain in the H3K27ac signal was primarily derived from the development of new enhancer elements. We next validated our ChIP-Seq results by performing ChIP-qPCR analysis of arbitrarily selected gained and lost regulatory elements in a new cohort of 16-day-old control and *Pkd1* mutant kidneys (**Suppl Fig. 2B**). Furthermore, to determine if our ChIP-Seq results can be extrapolated to other orthologous ADPKD models, we also performed H3K27ac ChIP using 21-day-old control and *Pkd2*-KO kidneys. We were able to validate *Pkd1*-mutant ChIP-Seq data even in *Pkd2*-KO kidneys, suggesting similar epigenetic rewiring in both ADPKD models (**Suppl Fig. 2C**).

**Figure 2.**
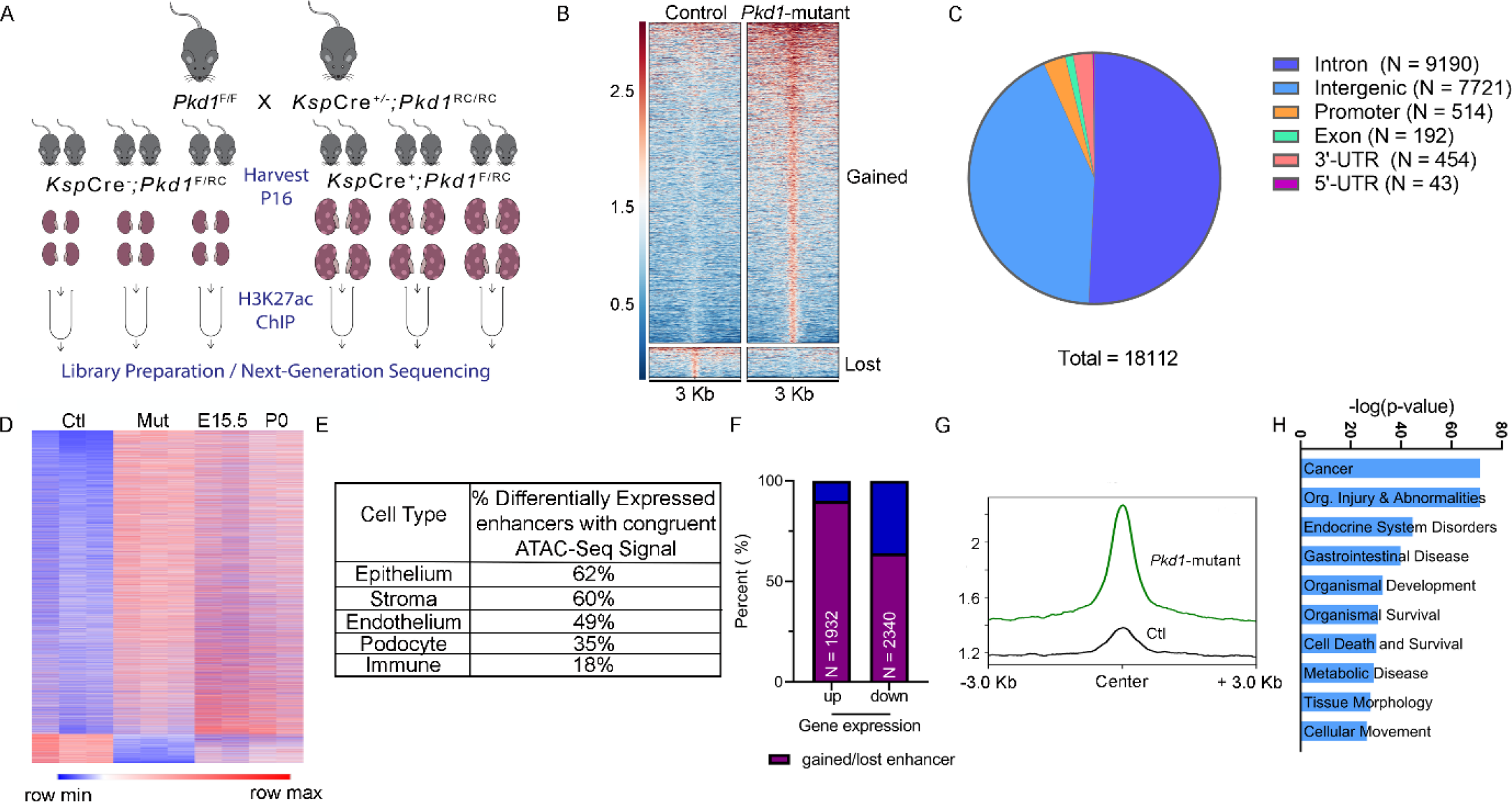
Comparative H3K27ac ChIP-seq uncovering the enhancer landscape in *Pkd1*- mutant kidneys. **A.** Graphic illustration of the comparative H3K27ac ChIP-Seq experimental design is shown. ChIP-seq was performed using three biological replicate samples, each containing pooled chromatin from four 16-day-old kidneys from control or *Pkd1*-mutant mice. **B.** Heatmap showing the signal intensity of H3K27ac (+/- 3kb) around the center of each differentially activated enhancer ordered by mean signal. *Pkd1-*mutant kidneys gained (higher H3K27ac level) 16560 enhancers and lost (lower H3K27ac level) 1552 enhancers. **C.** Pie chart depicting the genome-wide distribution of the differentially activated enhancers is shown. **D.** Heatmap shows the comparison of RPKM of differentially activated enhancers between control and *Pkd1*-mutant kidneys and wildtype E15.5 and P0 kidneys. **E.** The bulk H3K27Ac ChIP-seq data were deconvoluted using the E18.5 wildtype snATAC-seq dataset. The cellular distribution of activated enhancers is shown. **F.** Overlap of the *Pkd1-*mutant ChIP-Seq and RNA-Seq showing that >90% of upregulated genes are located in a TAD which co-houses an activated enhancer, whereas 64% of downregulated genes are located in a TAD which co-houses a lost enhancer. **G.** Average H3K27ac signal intensity of enhancer regions in control (black) and *Pkd1*-mutant (green) kidneys samples within TADs that houses upregulated genes. **H.** Ingenuity Pathway Analysis depicting the top pathways regulated by active enhancer upregulated gene pairs.

The transcriptomic profile of cystic kidneys has been likened to that of a developing kidney^46^. Therefore, we examined whether cystic and developing kidneys also share a similar epigenetic profile. We used ENCODE consortium H3K27ac ChIP-Seq datasets and mapped enhancers in embryonic (E) 15.5 and postnatal day (P) 0 kidneys^47, 48^. We then cross compared the ENCODE and our datasets and found that P16 non-cystic control kidneys have little resemblance to developing E15.5 or P0 kidneys. In contrast, we noted that *Pkd1*-mutant kidneys share 42% and 32% of H3K27ac peaks with E15.5 and P0 kidneys, respectively (**Fig. 2D and Suppl Fig. 3A-B**). Next, we deconvoluted our bulk H3K27ac ChIP-Seq data by integrating a recently published single-cell ATAC-seq dataset of wildtype mouse E18.5 kidney^41^. ATAC-seq is utilized to catalog genome-wide open chromatic positions, which is another indicator of active regulatory elements. In concordance with our findings of similarities with developing kidneys, 64% of active enhancers in 16-day-old *Pkd1*-mutant kidneys were found in regions with appropriately open/closed chromatin in the E18.5 kidneys (**Suppl Fig. 3C**). Moreover, this analysis revealed that our bulk H3K27ac signal primarily mapped to open chromatin locations in tubular epithelial and stromal cells (**Fig. 2E**).

**Figure 3.**
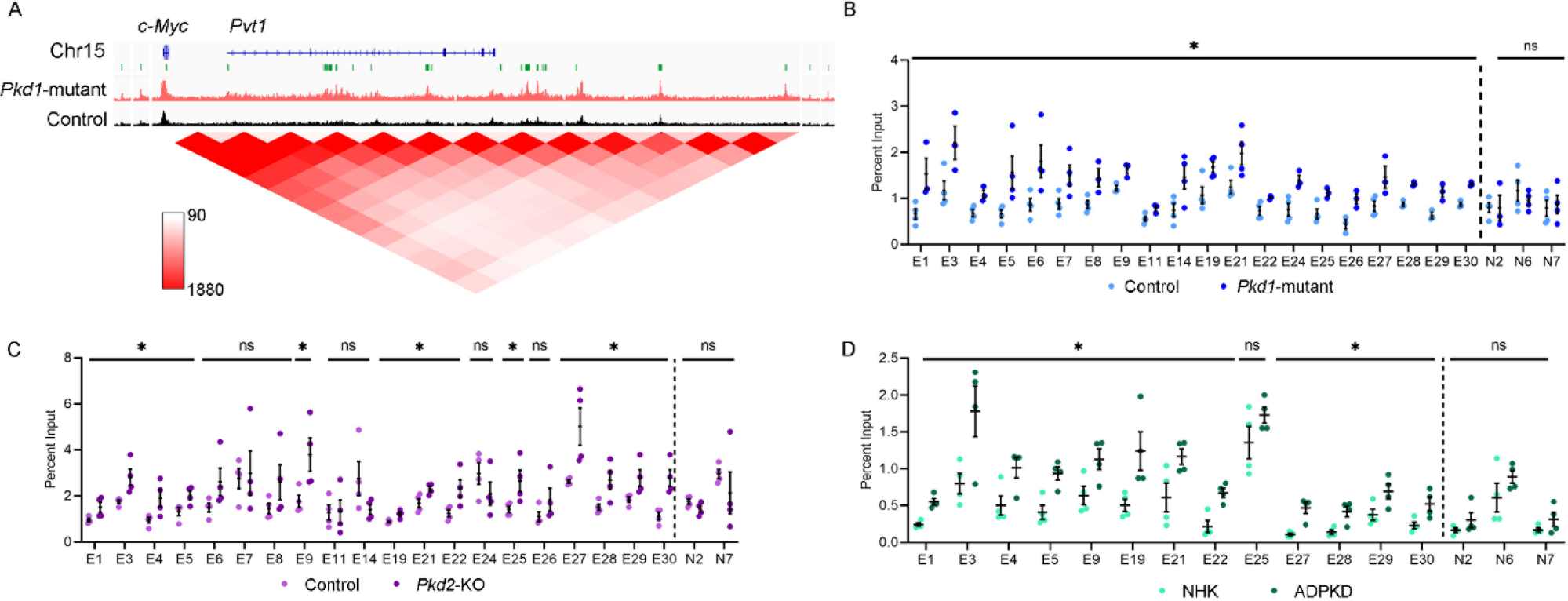
Evolutionarily conserved enhancer cluster in the *c-Myc* locus is activated in ADPKD models. **A.** ChIP-seq tracks showing a higher H3K27ac signal within the *c-Myc* locus in 16-day-old *Pkd1*-mutant compared to control kidneys. The activated enhancers are marked as green rectangles and denoted as E1-E30 (numbered in 5’ to 3’ orientation). A chromatin contact map (red) for the *c-Myc* locus derived from Hi-C mouse embryonic stem cell dataset is shown. **B-D.** ChIP-qPCR validation of the E1-E30 enhancers in *Pkd1*-mutant (B), *Pkd2*-KO (C), and human ADPKD kidneys (D) compared to their respective controls. N=3-4 all groups; error bars indicate SEM; * *P*<0.05. ns *P*>0.05; Student’s *t*-test.

Enhancers regulate gene expression by interacting with nearby or very distant promoters. However, topological associated domain (TAD) boundaries serve as physical limits on the range of enhancer-promoter interaction^42^. To determine the influence of enhancers on dysregulated gene expression in PKD, we first overlapped the H3K27ac ChIP-Seq data with our previously published RNA-seq dataset from 22-day-old control and *Pkd1*-mutant kidneys^21^. Next, we examined the relationship between active enhancers and dysregulated genes located within the same TAD boundaries. We found that 90% of upregulated genes in *Pkd1*-mutant kidneys were located in a TAD that also housed a gained enhancer (**Fig. 2F, G**). Conversely, 64% of downregulated genes were found in a TAD that had lost an enhancer. Finally, pathway analysis of the positively-correlated enhancer-dysregulated gene pairs revealed cancer signaling as the top influenced network (**Fig. 2H**). Closer examination of genes within the cancer signaling pathway identified *c-Myc* as the top gene with the most associated enhancers. Taken together, our analysis has uncovered a large repertoire of regulatory enhancers in cystic kidneys.

### Evolutionarily-conserved enhancer cluster transactivates c-Myc expression in cellular ADPKD model

The proto-oncogene *c-Myc* is transactivated in mouse and human ADPKD, and its overexpression is sufficient to induce kidney cysts in mice^21, 49, 50^. However, how c-Myc expression is regulated in PKD is incompletely understood. We identified 30 regulatory elements spanning 2.5 Mbp genomic region surrounding the *c-Myc* gene that displayed significantly enriched H3K27ac signal in *Pkd1*-mutant kidneys compared to control kidneys (**Fig. 3A**). A query of the publicly available high-throughput chromatin conformation capture (Hi-C) datasets revealed that these 30 active enhancers physically interact with the c-*Myc* promotor (**Fig. 3A**)^51^. We found that 22/30 enhancers are evolutionarily-conserved between mice and humans. Ten of these enhancers have not previously been reported in the literature (**Supplementary Table 1**)^52^. These observations suggest that the gained enhancers may underlie c-Myc upregulation in ADPKD and prompted us to further validate this locus. We began by performing ChIP-qPCR in an independent cohort (biological replicates) of 16-day-old control and *Pkd1*-mutant kidneys and confirmed that 20 conserved enhancers indeed display higher H3K27ac levels in cystic kidneys (**Fig. 3B**). As a pertinent negative control, we found several regions that exhibit equal H3K27ac signal in our ChIP-Seq data remained unchanged by ChIP-qPCR validation in control and *Pkd1*-mutant kidneys (**Fig. 3B**). Next, we asked which of these enhancers were also enriched in *Pkd2*-KO kidneys. ChIP-qPCR revealed that 13/20 enhancers have higher H3K27ac signal in 21-day-old *Pkd2*-KO kidneys compared to age-matched control kidneys (**Fig. 3C**). Finally, we examined whether these enhancers are relevant to human ADPKD. ChIP-qPCR showed that 12/20 enhancers exhibited higher H3K27ac modification in the human ADPKD samples than NHK control samples (**Fig. 3D**). Taken together, our careful assessment of the *c-Myc* locus identified 12 evolutionarily-conserved enhancers that are activated in both mouse and human ADPKD samples.

To determine if these enhancers drive c-Myc expression, we used CRISPR/Cas9 to delete them in a cellular *Pkd1*^RC/-^ ADPKD model. The *Pkd1*^RC/-^ cells are of collecting duct origin (see methods for details) that harbor a hypomorphic p.R3277C mutation on one *Pkd1* allele, whereas the other is deleted^53^. Isogenic *Pkd1*^RC/+^ cells serve as controls. Our characterization of this new cellular model revealed that it recapitulates the key pathogenic hallmarks of *Pkd1*-mutant mice. We found that *Pkd1^RC/-^* cells have reduced Polycystin-1 (but not Polycystin-2) expression (**Fig. 4A**) and higher 3D cyst growth (**Fig. 4B**), and proliferation rates (**Fig. 4C**) than *Pkd1^RC/+^* cells. Importantly, we noted activated pathogenic ADPKD signaling, including higher c-Myc, Fos, and CREB expression in *Pkd1^RC/-^* compared to *Pkd1^RC/+^* cells (**Fig. 4A and Suppl. Fig 4**). Moreover, as is the case with mouse and human ADPKD samples, ELISA revealed 35% higher total H3K27ac levels in *Pkd1*^RC/-^ cells compared to *Pkd1*^RC/+^ cells (**Fig. 4D**). Moreover, ChIP-qPCR showed increased H3K27ac modification on 10/22 conserved c-Myc-flanking enhancers in *Pkd1*^RC/-^ cells compared to *Pkd1*^RC/+^ cells (**Fig. 4E**).

**Figure 4.**
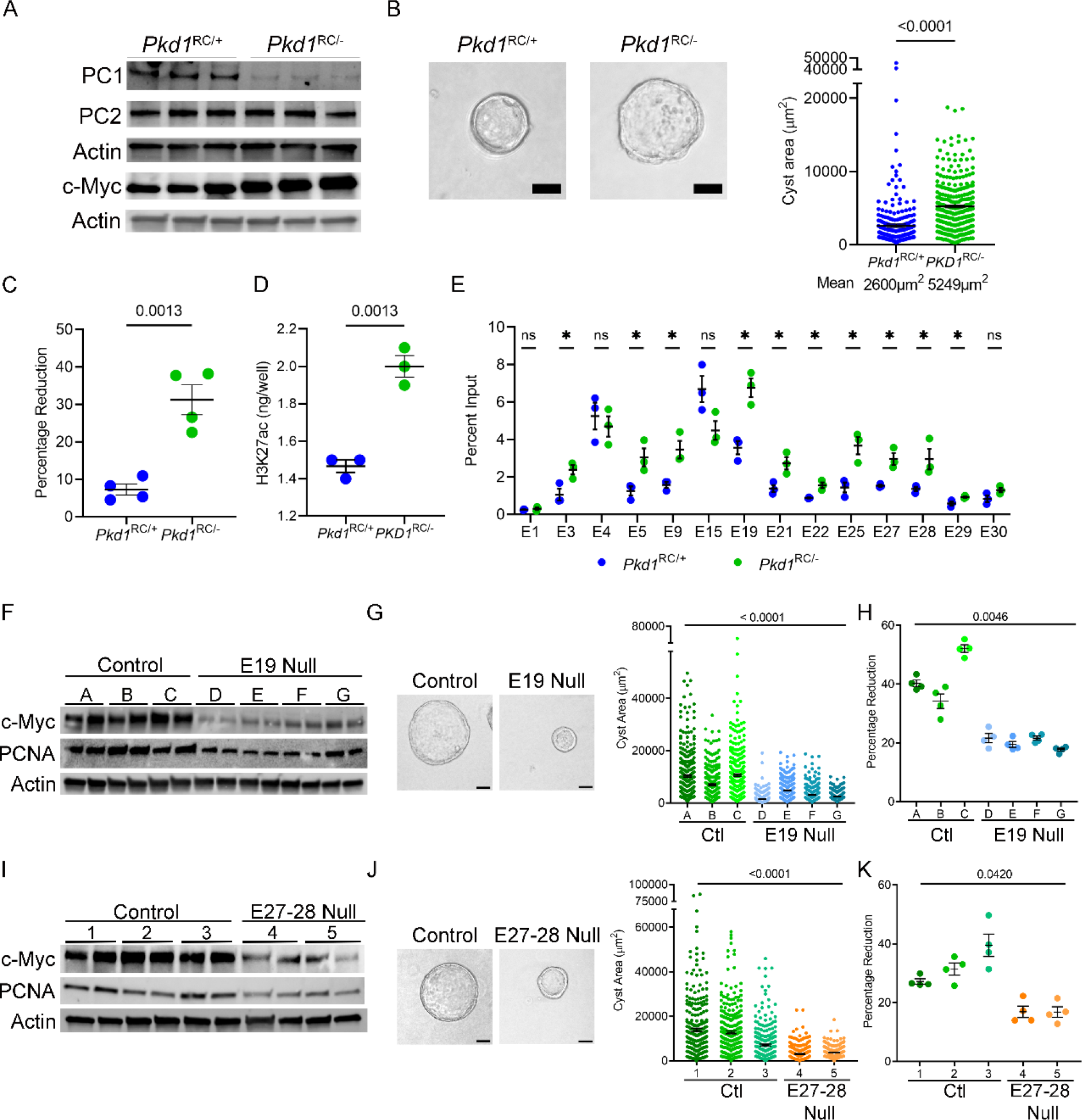
Enhancers activate c-Myc and regulate proliferation and cyst growth in *Pkd1*- mutant cells. **A.** Western blot analysis showing reduced Polycystin1 (PC1) expression in *Pkd1*^RC/-^ cells compared to parental *Pkd1*^RC/+^ cells. Polycystin2 (PC2) expression remained unchanged, whereas c-Myc expression was increased in *Pkd1*^RC/-^ compared to *Pkd1*^RC/+^ cells. Actin serves as the normalizing loading control. **B.** Representative images and quantification showing increased cyst size of *Pkd1*^RC/-^ compared to *Pkd1*^RC/+^ cells grown in 3D Matrigel for 7- days. **C.** Alamar Blue measurement 12 hours after incubation showing increased proliferation of *Pkd1*^RC/-^ compared to *Pkd1*^RC/+^cells. **D.** ELISA showing higher global H3K27ac levels in *Pkd1*^RC/-^ compared to *Pkd1*^RC/+^ cells. **E.** Comparative H3K27ac ChIP-qPCR validation of the *c-Myc* locus enhancers in *Pkd1*^RC/-^ and *Pkd1*^RC/+^ cells. **F.** Western blot analysis showing reduced c-Myc expression in *Pkd1*^RC/-^ cell lines lacking the E19 enhancer compared to the unedited *Pkd1*^RC/-^parental cells. **G.** Representative images and quantification showing reduced cyst size of E19- edited compared to unedited *Pkd1*^RC/-^ cell lines grown 3D Matrigel for 7-days. **H.** Alamar Blue- assessed proliferation rate of E19-edited and unedited *Pkd1*^RC/-^ cells is shown. **I-K**. Western blot analysis, images and quantification of 3D cyst size, and Alamar Blue measurements showing reduced c-Myc expression, cyst size, and proliferation of *Pkd1*^RC/-^ cells lacking the E27-28 enhancer cluster compared to the unedited parental *Pkd1*^RC/-^ cells. Images were taken at 20x magnification. N=300 (from 3 biological replicates) for all cyst measurements. N=4 biological replicates for all Alamar blue measurements. Error bars indicate SEM. * *P* < 0.05. ns = *P* > 0.05. Statistical analysis: Students *t-*test (B-E) and nested *t*-test (G, H, J, K). Scale bars represent 25 μM.

The enhancer elements are referred to as E1 through E30, labeled based on their genomic position (in 5’-3’ orientation) within the TAD, which houses the *c-Myc* gene. First, we deleted the 3 kb region, encompassing the E19 enhancer in *Pkd1^RC/-^* cells. We generated four independent clonal cell lines lacking the E19 enhancer, and in all four cell lines, we noted a reduction in c-Myc expression (**Fig. 4F, Suppl. Fig. 5A**). We also noted that 3D cyst growth was reduced by 50% in E19 null clones compared to their respective unedited *Pkd1*^RC/-^ parental clones (**Fig. 4G**). Moreover, cellular proliferation as measured by the Alamar Blue assay was reduced, and western blot analysis revealed lower PCNA levels in each of the E19 null clones compared to unedited clones (**Fig. 4H**). Next, we generated two clonal cell lines that lacked a 1.6 kb region encompassing E27 and E28 enhancers **(Suppl Fig 5B)**. Again, we noted reduced c-Myc and

**Figure 5.**
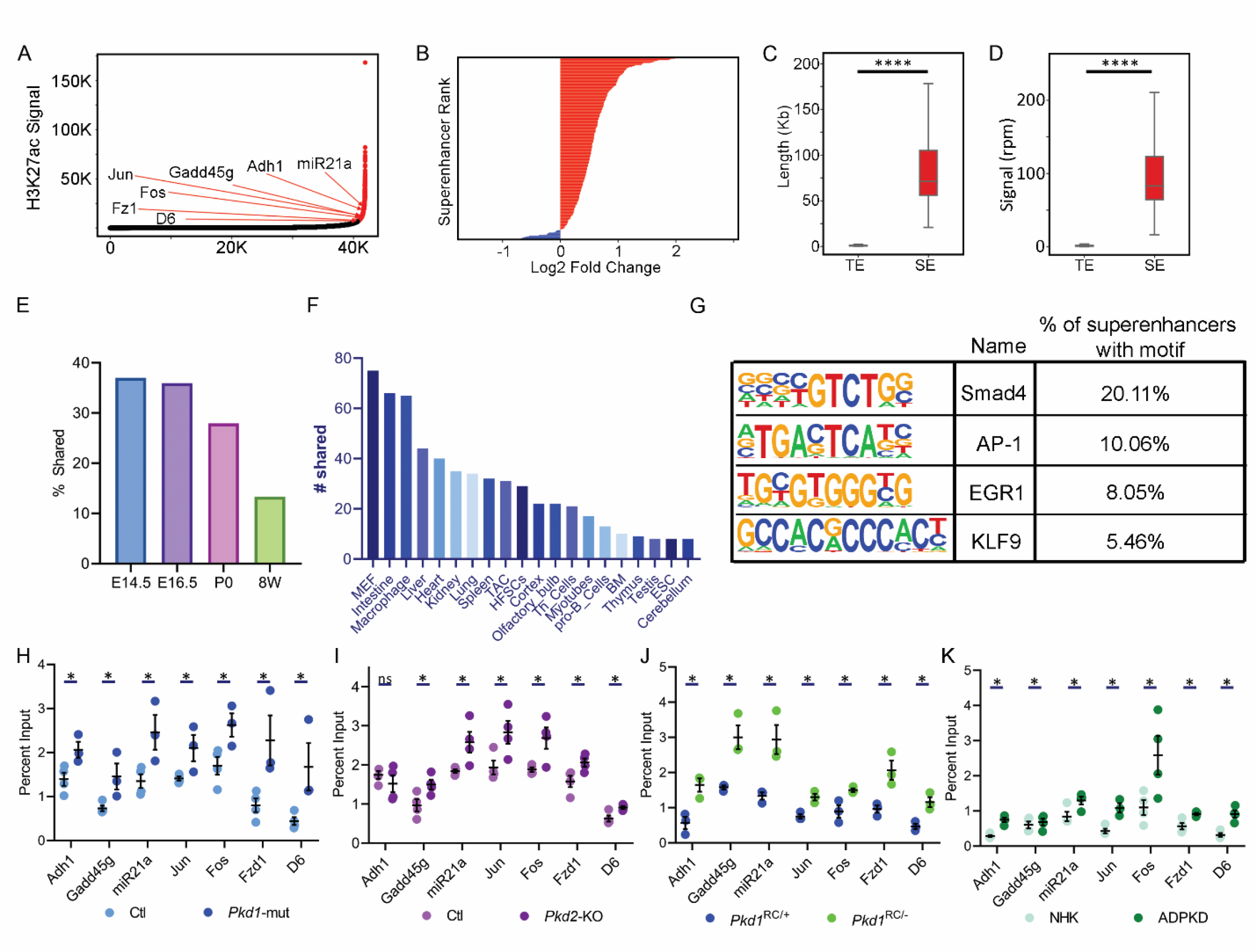
Super-enhancer landscape of ADPKD. **A.** Hockey stick plot showing rank-ordered, input normalized H3K27ac signal in *Pkd1*-mutant and control kidneys. The CREs meeting the super-enhancer criteria are highlighted in red. Selected super-enhancers annotated based on nearest neighboring genes are shown. **B.** The graph depicts all differentially activated super-enhancers rank-ordered based on Log2 fold change in the H3K27ac signal in 16-day-old *Pkd1*- mutant compared to control kidneys. Red indicates super-enhancers with higher and blue indicates those with lower H3K27ac signal in *Pkd1*-mutant compared to control kidneys. One hundred five super-enhancers were gained, and five were lost in *Pkd1*-mutant compared to control kidneys. **C and D.** Quantification of genomic length and H3K27ac signal intensity of super-enhancers (SE) versus total enhancers (TE) is shown. **E.** Super-enhancers of wildtype E14.5, E16.5, P0, and 8-week (W) kidneys were mapped using the ENCODE data. The graph depicts the level of overlap in the super-enhancer landscape of cystic *Pkd1*-mutant kidneys and normal developing and adult kidneys. **F.** The *Pkd1*-mutant and dbSUPER super-enhancer datasets were cross-compared. The graph depicts the common super-enhancers found in both databases broken down based on cell and tissue type. **G.** Motif analysis of super-enhancers using the Homer software is shown. **H-K.** ChIP-qPCR validation of selected super-enhancers in *Pkd1*-mutant and *Pkd2*-KO kidneys, *Pkd1*^RC/-^ cells, and human ADPKD samples compared to their respective controls. N = 3-4 all groups. * *P* < 0.05, **** *P* <0.001. ns = *P* > 0.05. Error bars indicate SEM. Statistical analysis: Nested *t*-test (C and D), Student’s *t*-test (H-K)

PCNA expression (**Fig. 4I**), a 30% reduction in cyst size (**Fig. 4J**), and a lower proliferation rate (**Fig. 4K**) in E27-28 deleted *Pkd1*^RC/-^ cell lines compared to their unedited counterparts. We also deleted the genomic locus containing E5-E9 enhancers (**Suppl Fig. 6A**). However, this deletion was not sufficient to reduce *Myc*. Finally, we attempted to delete three additional loci: 4.9kb genomic region containing E21-22, 868 bp genomic region containing E25, and 498bp genomic region containing E29. However, we could only recover heterozygous deletions of E21-22 and E25 (**Suppl Figure 6B,C)**. We did not observe differences in c-Myc expression in any of the heterozygous deletion lines. Similarly, we did not obtain even heterozygous deletion of the genomic region encompassing E29 despite multiple attempts. Thus, genomic loci bearing E19, E27, and E28 enhancers promote c-Myc expression in cellular ADPKD models. **Super-enhancers promote Fos and Jun expression**: Super-enhancers are large genomic regions that contain multiple enhancers in close proximity to each other. We used the ROSE (Rank Ordering of Super-Enhancers) bioinformatic tool to map ADPKD-relevant super-enhancers from our H3K27ac ChIP-Seq dataset^17, 37^. Using differential binding analysis, we found that *Pkd1*- mutant kidneys gained 101 (higher H3K27ac) and lost 5 (lower H3K27ac) super-enhancers compared to control kidneys (**Fig. 5A-B**). These super-enhancers, on average, were 71 kb in genomic length and exhibited greater than 50-fold higher H3K27ac modification compared to a typical enhancer (**Fig 5C and D)**. We also mapped the super-enhancer landscape of E14.5, E16.5, and P0 developing kidneys using the ENCODE data. As with enhancers, we found that many *Pkd1*-mutant gained super-enhancers were also active in developing kidneys (**Fig. 5E**). Interestingly, cross-comparison with the dbSUPER super-enhancer database revealed that numerous gained super-enhancers were enriched in stromal and immune cells, suggesting epigenetic rewiring even in the cyst microenvironment (**Fig. 5F**). Finally, motif analysis revealed that the AP-1 binding element was significantly enriched amongst the gained super-enhancers (**Fig 5G**).

**Figure 6.**
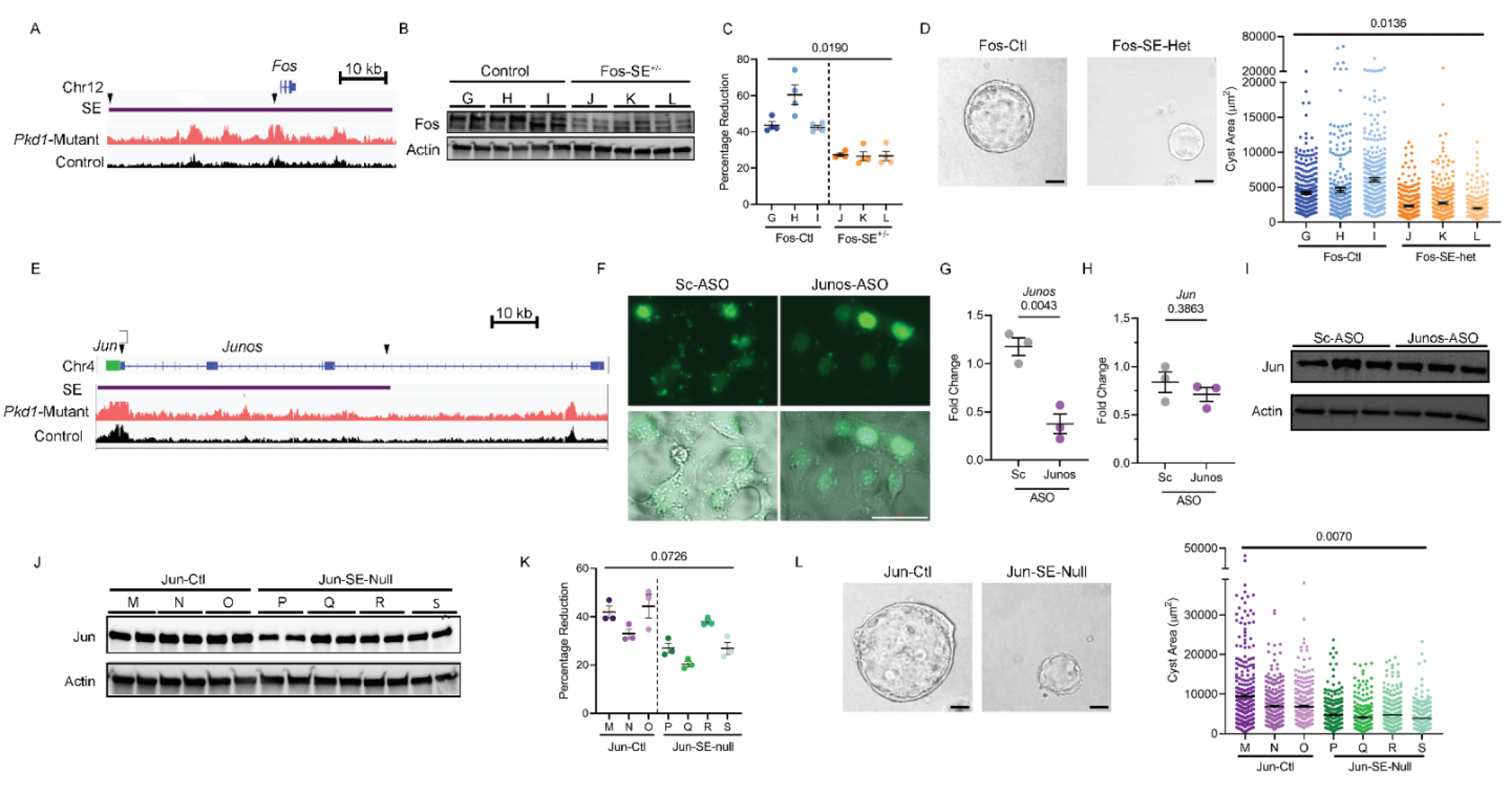
Super-enhancers activate Fos and Jun and regulate proliferation and cyst growth in *Pkd1*-mutant cells. **A.** ChIP-seq tracks of the genomic locus encompassing the *Fos* gene showing higher H3K27ac modification in *Pkd1*-mutant compared to control kidneys. The Fos super-enhancer is denoted by the purple rectangle. Arrows indicate sites of sgRNAs for CRISPR/Cas9 mediated deletion of the Fos super-enhancer. **B.** Western blot analysis showing reduced Fos abundance *Pkd1*^RC/-^ cells with heterozygous deletion of the Fos super-enhancer (Fos-SE^+/-^) compared to the unedited parental cell lines (Control). **C.** Alamar Blue-assessed proliferation of control and Fos-SE^+/-^ cell lines is shown. **D.** Representative images and quantification showing reduced cyst size in Fos-SE^+/-^ cell lines compared to control cell lines. **E.** ChIP-seq tracks of the *Jun* genomic locus showing higher H3K27ac modification in *Pkd1*-mutant compared to control kidneys. *Jun* gene is denoted by green rectangle, lncRNA *Junos* is denoted by blue, and the super-enhancer is denoted by a purple rectangle. Arrows indicate sgRNA targeted sites for super-enhancer deletion. **F.** *Pkd1*^RC/-^ cells were transfected with fluorescein amidite (FAM) labeled control antisense oligonucleotide (ASO) or an ASO targeting *Junos* lncRNA. Fluorescent and bright-field microscopic images showing delivery of control and *Junos* ASOs into cells 24-hours after transfection. **G.** qPCR analysis demonstrated reduced *Junos* transcript abundance in Junos ASO-treated compared to control ASO-treated *Pkd1*^RC/-^ cells. **H and I.** qPCR and western blot analysis showing that Jun expression was not different in *Pkd1*^RC/-^ cells treated with control compared to *Junos* ASO. **J.** Western blot analysis showing reduced Jun expression in *Pkd1*^RC/-^ cell lines lacking the 62kb super-enhancer (Jun-SE-null) compared to parental unedited *Pkd1*^RC/-^ cells (Jun-ctl). **K.** Alamar Blue assay demonstrating reduced proliferation in Jun-SE-null compared to Jun-ctl *Pkd1*^RC/-^ cells. **L.** Representative images and quantification showing reduced cyst size in Jun-SE-null compared to Jun-ctl cells. Error bars indicate SEM; Statistical analysis: Nested *t*-test (C, D, K, and L) and Student’s *t*-test (G and H). Scale bars represent 25μM

AP-1 is a tetramer transcription factor comprised of *Jun* and *Fos*. Interestingly, we noted that *Jun* and *Fos* loci themselves are flanked by gained super-enhancers. Considering both Jun and Fos are implicated in mouse and human ADPKD, we asked whether these super-enhancers are relevant to cyst growth. First, ChIP-qPCR in control and *Pkd1*-mutant kidneys confirmed higher H3K27ac modification on Fos, Jun, and a series of other randomly selected super-enhancers (**Fig. 5H**). Moreover, we found this super-enhancer landscape was also observed in *Pkd2*-mutant kidneys, *Pkd1*^RC/-^ cells, and human ADPKD samples compared to their respective controls (**Fig. 5I-K**). The Fos super-enhancer encompasses a 35 kb region upstream of the *Fos* gene promoter (**Fig. 6A**). We used CRISPR/Cas9 to delete the upstream super-enhancer while leaving the Fos promoter intact in *Pkd1*^RC/-^ cells. Compared to unedited *Pkd1*^RC/-^ cells, we found that *Pkd1*^RC/-^ cells with a heterozygous deletion of the Fos super-enhancer had reduced Fos expression (**Fig. 6B**). Moreover, the edited *Pkd1*^RC/-^ cells had lower cellular proliferation and cyst growth rates compared to the unedited *Pkd1*^RC/-^ cells (**Fig. 6C and D**).

The putative Jun super-enhancer is an uncharacterized 62 kb genomic region upstream of the *Jun* gene. Closer examination revealed that this super-enhancer region also encodes a long noncoding RNA (lncRNA) annotated as *Junos* (**Fig. 6E**). Therefore, the locus may act as a *Jun* super-enhancer, or alternatively, it could regulate Jun expression via *Junos*. To determine whether the lncRNA regulates *Jun* expression, we inhibited *Junos* in *Pkd1*^RC/-^ cells using antisense oligos (ASOs) while still keeping the super-enhancer intact (**Fig. 6F**). ASO treatment reduced *Junos* level by 68 percent (**Fig. 6G**). However, *Jun* expression remained unchanged, indicating that the lncRNA does not regulate *Jun* expression (**Fig. 6H-I**). Next, we used CRISPR/Cas9 and generated three clonal cell lines with homozygous deletion of the 70 kb super-enhancer region. Compared to the unedited *Pkd1^RC/-^* control cells, Jun expression was reduced in all three *Pkd1^RC/-^* cell lines lacking the super-enhancer (**Fig. 6J**). Moreover, cellular proliferation and cyst size were also significantly reduced in cell lines Jun super-enhancer deletion (**Fig. 6K-L**). Thus, both components of the AP-1 complex are trans-activated by neighboring super-enhancers in *Pkd1*- mutant cells.

### ADPKD metabolic pathways influence H3K27ac levels

Histone acetylation is a dynamic process tied to acetyl-CoA availability, which is controlled by the cellular metabolic state^54–56^. ADPKD is marked by extensive metabolic reprogramming, including the activation of aerobic glycolysis and glutaminolysis and inhibition of oxidative phosphorylation. Therefore, we asked whether the ADPKD metabolic pathways regulate H3K27ac levels. First, we treated *Pkd1*^RC/-^ cells with 2-deoxyglucose (2-DG) for 24 hours and then measured H3K27ac by ELISA. 2-DG is a competitive inhibitor of aerobic glycolysis. We noted that 2-DG treatment in *Pkd1*^RC/-^ cells reduced H3K27ac levels by 30% (**Fig. 7A**). Moreover, H3K27ac modification was reduced at c-Myc enhancers with 2-DG treatment in *Pkd1*^RC/-^ cells (**Fig. 7B**). Accordingly, c-Myc expression was reduced in 2-DG treated *Pkd1*^RC/-^ cells compared to untreated *Pkd1*^RC/-^ cells (**Fig. 7C**). Next, we treated *Pkd1*^RC/-^ cells with BPTES, an allosteric inhibitor of the glutaminase enzyme, to turn off glutaminolysis or WY-14643, a *Ppara* agonist, to activate oxidative phosphorylation. We noted that compared to untreated *Pkd1*^RC/-^ cells, total H3K27ac levels and H3K27ac abundance at c-Myc enhancers was also reduced in *Pkd1*^RC/-^ cells treated with BPTES or WY-14643 (**Fig 7D, E, G, and H)**. Accordingly, Western blot analysis revealed that treatment with both BPTES and WY-14643 reduced *c-Myc* expression (**Fig 6F and I**). Thus, the rewiring of the epigenetic profile is at least in part mediated by ADPKD metabolic reprogramming.

**Figure 7.**
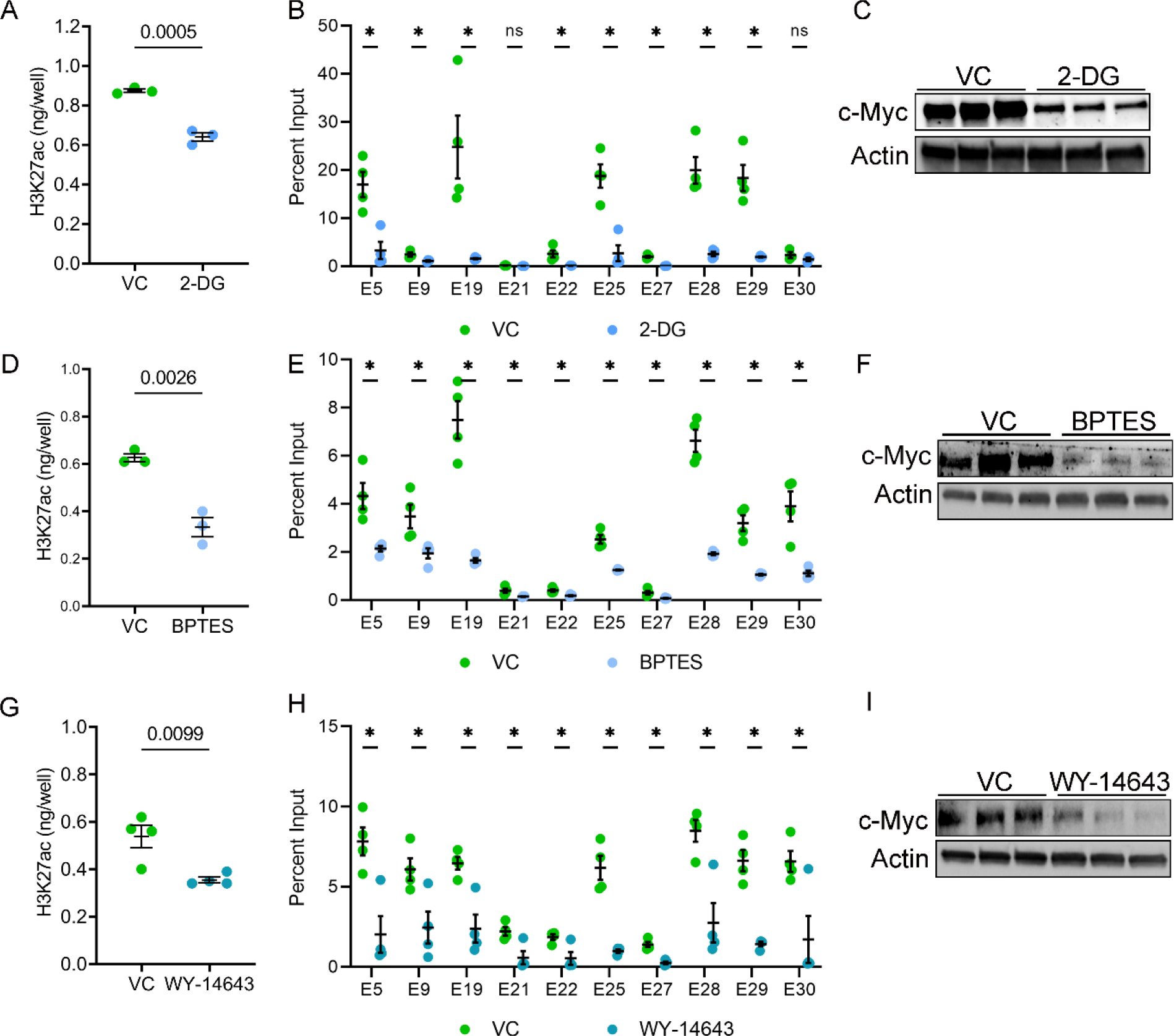
Metabolic pathways influence H3K27ac levels in *Pkd1*-mutant cells. **A.** *Pkd1*^RC/-^ cells were treated with 2-DG, an inhibitor of aerobic glycolysis, or vehicle control for 24 hours. ELISA revealed that histone extracts from 2-DG-treated cells had reduced H3K27ac levels compared to vehicle-treated cells. **B.** ChIP-qPCR showed reduced H3K27ac modification on *c-Myc* enhancers in *Pkd1*^RC/-^ cells treated with 2-DG compared to vehicle control. **C.** Western blot analysis showing reduced c-Myc expression in 2-DG-treated compared to vehicle-treated *Pkd1*^RC/-^ cells. **D.** *Pkd1*^RC/-^ cells were treated for 48 hours with vehicle or BPTES to inhibit glutaminolysis. ELISA revealed that BPTES-treated cells had a reduced level of H3K27ac histone modification compared to vehicle-treated cells. **E.** ChIP-qPCR showing reduced H3K27ac modification on c-Myc locus enhancers in BPTES-treated compared to vehicle-treated *Pkd1*^RC/-^ cells. **F.** Western blot analysis showing reduced c-Myc expression in *Pkd1*^RC/-^ cells treated with BPTES compared to control vehicle. **G-I.** *Pkd1*^RC/-^ cells were treated for 72 hours with the Ppara agonist WY-4643 or control vehicle. ELISA revealed reduced global H3K27ac levels, ChIP-qPCR showed lower H3K27ac signal on c-Myc enhancers, and Western blot analysis demonstrated c-Myc downregulation in WY-4643-treated compared to vehicle-treated *Pkd1*^RC/-^ cells. N=3-4 all groups. * *P* < 0.05, ns = *P* >0.05. Error bars indicate SEM. Statistical analysis: Student’s *t*-test.

## DISCUSSION

Large-scale transcriptomic rewiring is a prominent pathological feature of ADPKD. Here, we provide an in-depth view of the cystic kidney enhancer landscape that underlies this dysregulated gene expression. The first novel insight of our work is that there is a marked increase in global H3K27ac levels in mouse and human ADPKD tissues. Further, our findings suggest this increase, at least in part, is a consequence of ADPKD metabolic reprogramming involving aerobic glycolysis and glutaminolysis. Indeed, both metabolic pathways are known to replenish and exaggerate the cellular acetyl-CoA pool, which in turn is the critical determinant for histone acetylation^54–56^. Conversely, work from others suggests that super-enhancers propagate the aberrant expression of metabolic pathway genes in *Pkd1*-mutant cells ^57^. Thus, there may be a vicious metabolism- epigenetics cycle in ADPKD, where the altered metabolism reshapes the epigenetic state of cyst epithelia, leading to a transcriptomic output conducive to sustained metabolic reprogramming.

The second and perhaps the most important outcome of our studies is identifying the genome-wide enhancer profile of *Pkd1*-mutant kidneys. We uncovered >16,000 preferentially activated CRE, including 105 super-enhancers, providing the first glimpse of extensive epigenetic rewiring of cystic kidneys. Some noteworthy observations were that: (i) the activated CREs are located in intergenic as well as intronic regions, implying both long and short-range enhancer-promoter looping events in PKD. (ii) Deconvolution of our global H3K27ac ChIP-seq data using E18.5 wildtype kidney scATAC-seq data suggested that activated enhancers are observed both in cyst epithelia and cyst microenvironment component stromal and immune cells. (iii) We noted that >90% of upregulated genes are located in TADs that also housed activated enhancers suggesting that majority of the dysregulated transcriptome is facilitated by the altered epigenetic landscape.

c-Myc and the AP-1 heterodimer complex components Fos and Jun are transactivated in ADPKD models ^22^. In turn, these transcription factors activate pro-proliferative gene networks that are thought to underlie cyst growth ^58, 59^. Our work provides several new mechanistic insights into the regulation of this transcriptional circuitry. Our data suggest that a series of intergenic enhancers are necessary to drive c-Myc upregulation in ADPKD. Consistent with these observations, enhancer-mediated upregulation of c-Myc has also been reported in malignancies^52^, albeit the enhancer choice appears to vary. Interestingly, 10/12 activated enhancers in both mouse and human ADPKD samples have not been reported in cancers, suggesting that c-Myc enhancer repertoire and choice may differ based on the type of tissue and disease. In support of this notion, deletion of a large enhancer 1.7 megabases downstream of the *c-Myc* gene abolishes c-Myc expression but only in hematopoietic stem cells and appears to be critical for leukemic transformation ^60^. Therefore, it is tempting to speculate that the 10 ADPKD- enriched enhancers double as regulators of c-Myc expression during kidney development. Moreover, these key enhancers could serve as novel targets to reduce c-Myc expression preferentially in the kidney and slow cyst growth.

Thematically similar to the c-Myc locus, we found that super-enhancer elements are necessary to upregulate Jun and Fos in *Pkd1*-mutant cells. The same Fos super-enhancer is also functional in neuronal cells suggesting that this CRE is active in diverse cell types ^61^. In contrast, little was known about the 62 kb super-enhancer downstream of *Jun*. We report that this locus serves dual functions of a *Jun* super-enhancer and *Junos* lncRNA gene. Super-enhancer-associated lncRNAs are implicated in cis-activating nearby genes via mechanisms such as assisting in chromatin looping or transcription factor binding. Instead, we found that *Junos* was dispensable for Jun expression, implying an alternative biological role of this lncRNA, perhaps in trans-activating other target genes.

There are limitations and caveats related to our work. First, TAD-specific active enhancer and upregulated gene pairs indicate positive correlation but do not imply a direct causal link. While we established causation for c-Myc, Fos, and Jun, all other enhancer-gene pairs remain unvalidated. High throughput CRISPR-based and massively parallel reporter assays have been recently described to study regulatory functions of enhancers^62^. Our dataset will be a valuable resource to apply these approaches. Second, integrating our H3K27ac ChIP-seq dataset with the Hi-C dataset provides a good view of the extensive chromatin looping in *Pkd1*-mutant kidneys. However, an independent Hi-C dataset for cystic kidneys is needed for a refined and accurate description of the 3D chromatin architecture in PKD. Third, we did not validate c-Myc, Jun, or Fos CREs in ADPKD mouse models. These studies will require new mutant mouse models and are the focus of our future work. Finally, we observed congruent epigenetic signatures across the various ADPKD models and human samples based on ChIP-qPCR of arbitrarily selected enhancers. However, independent genome-wide datasets need to be developed in other ADPKD models, including human ADPKD tissues.

In summary, we provide the first insights into the enhancer map of *Pkd1*-mutant kidneys. We identify a series of enhancers near pro-proliferative transcription factors and demonstrate their ability to regulate cyst growth and gene expression. Our work also suggests that H3K27ac levels are regulated by rewired metabolism of *Pkd1*-mutant cells. Finally, developing pharmacological approaches to inhibit these enhancers could serve as a novel method to slow cyst growth.

## Disclosures

None

## Acknowledgments

This work is dedicated to the memory of Ms. Peyton Johnson, who we lost in 2020. We are grateful for her technical and intellectual contributions in this project. We thank Chun-Mien Chang for help with 3D cystogenesis, the PKD Research and Biomaterials and Cellular Models Core at the University of Kansas Medical Center for human tissue samples, and the McDermott Center Next Generation Sequencing Core and Bioinformatics Core Facility (BICF) at UT Southwestern Medical Center for providing critical services.

Work from the authors’ laboratory is supported by National Institute of Diabetes and Digestive and Kidney Diseases Grants K08DK117049 (to R.L) and R01 DK102572 (to V.P.). R. L. is also supported by grants from the PKD Foundation and American Society of Nephrology KidneyCure Grants Program. V.M. is supported by the Cancer Prevention and Research Institute of Texas (RP150596)

## SUPPLEMENTARY INFORMATION

**Supplementary Figure 1.**
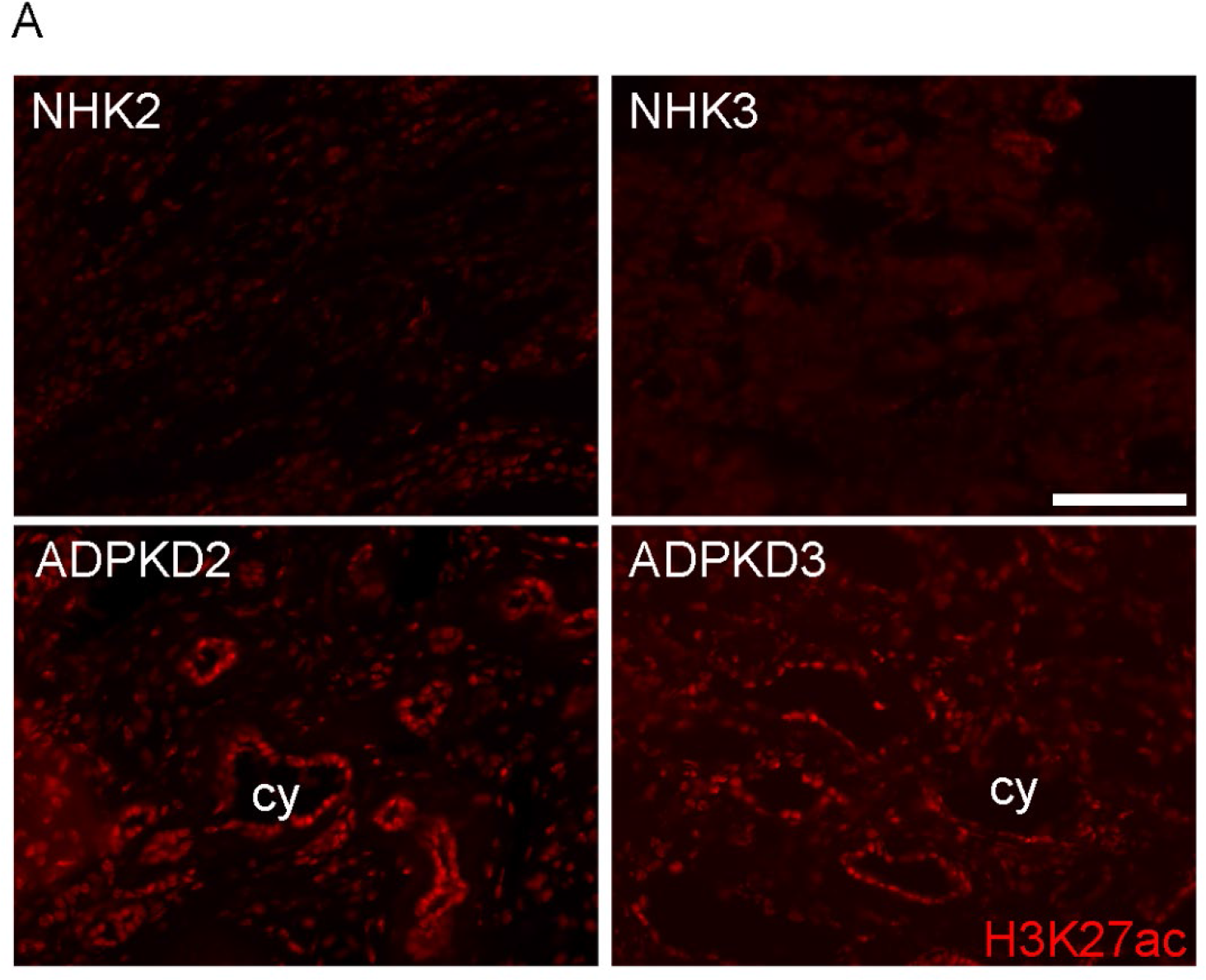
Immunofluorescence images showing anti-H3K27ac antibody staining of sections from two normal human kidney (NHK) and human ADPKD samples. Scale bar represents 100 μM. Cy = cyst

**Supplementary Figure 2:**
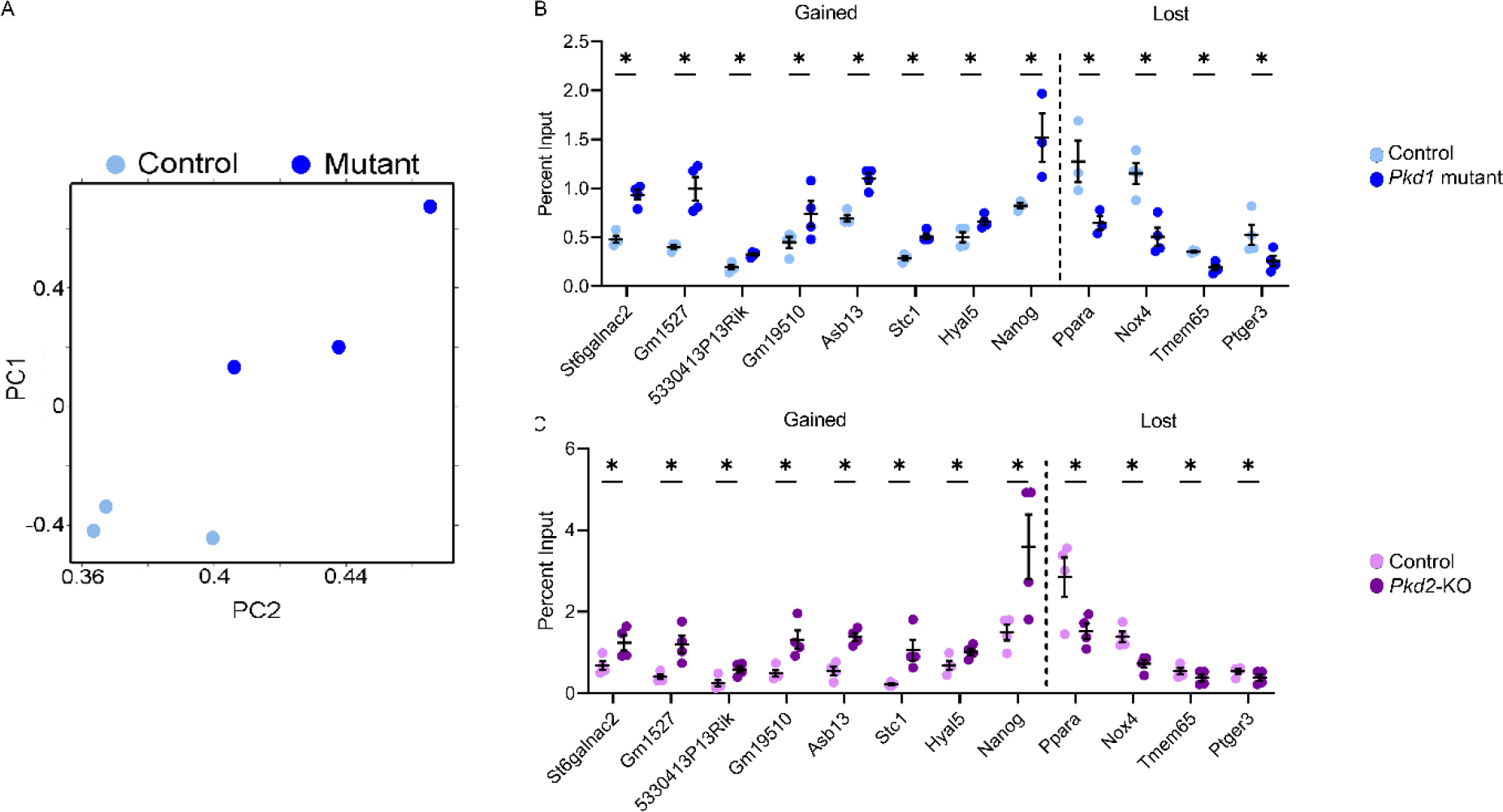
**A.** Principal component analysis of ChIP-Seq data from 16-day-old control and *Pkd1-*mutant kidneys. **B and C.** ChIP-qPCR validation of gained and lost enhancers in *Pkd1*-mutant and *Pkd2*-KO mice kidneys compared to age-matched littermate controls. N=3-4; * *P*<0.05; error bars represent SEM. Statistical analysis: Student’s t-test.

**Supplementary Figure 3.**
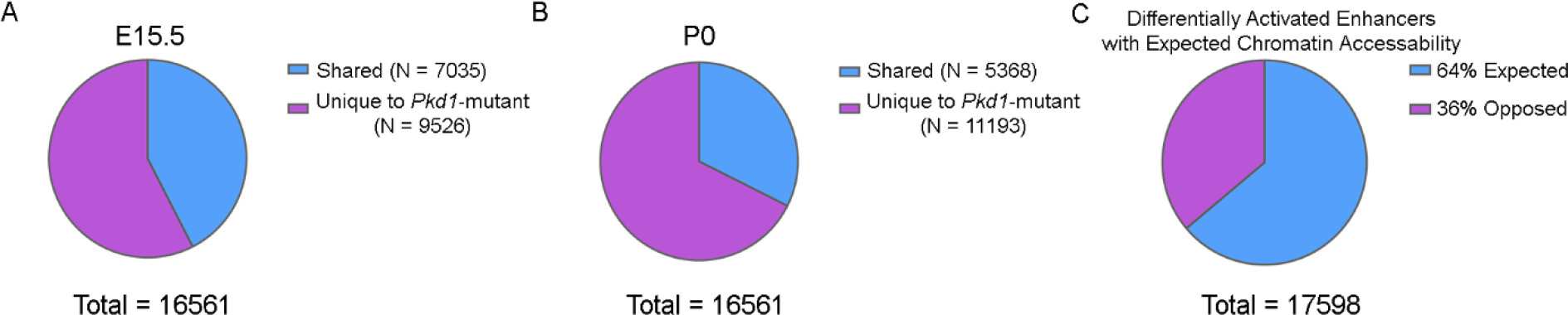
**(A and B)** Pie chart demonstrating overlap of the *Pkd1*-mutant enriched enhancers with those found in E15.5 and P0 kidneys. (**C)** Pie chart showing the percentage of differentially activated enhancers that fall into areas with expected chromatin accessibility.

**Supplementary Figure 4.**
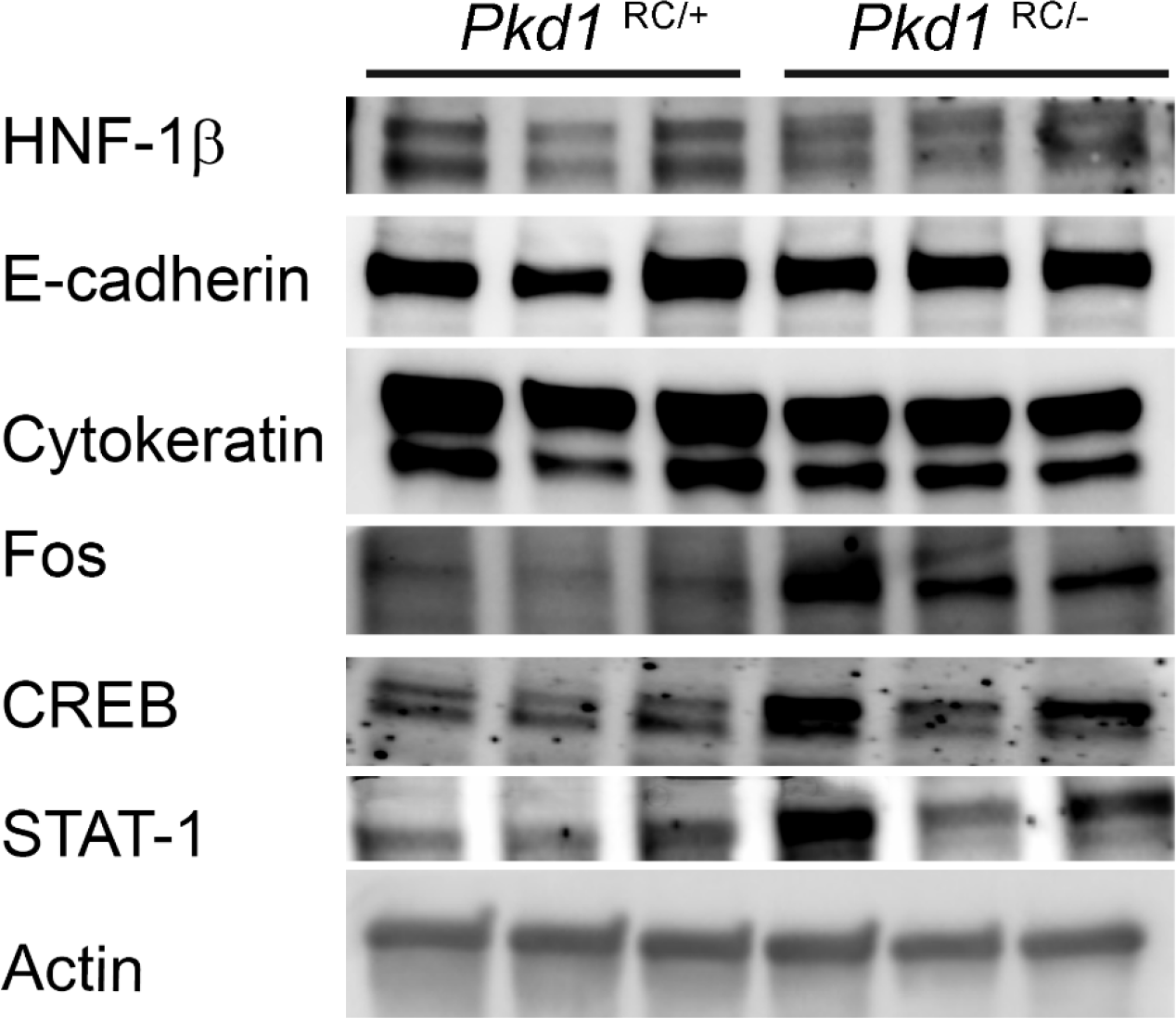
Western blot analysis of indicated proteins in *Pkd1*^RC/+^ and *Pkd1*^RC/-^ cell lines.

**Supplementary Figure 5.**
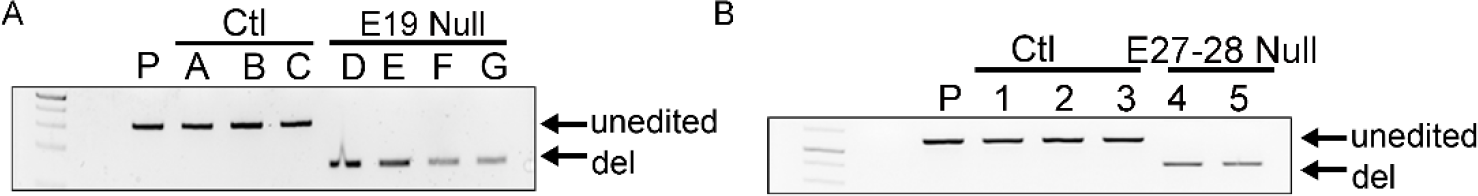
A, B. Genotyping confirmation of E19 and E27-28 null cell lines. P = parent control cell line.

**Supplementary Figure 6.**
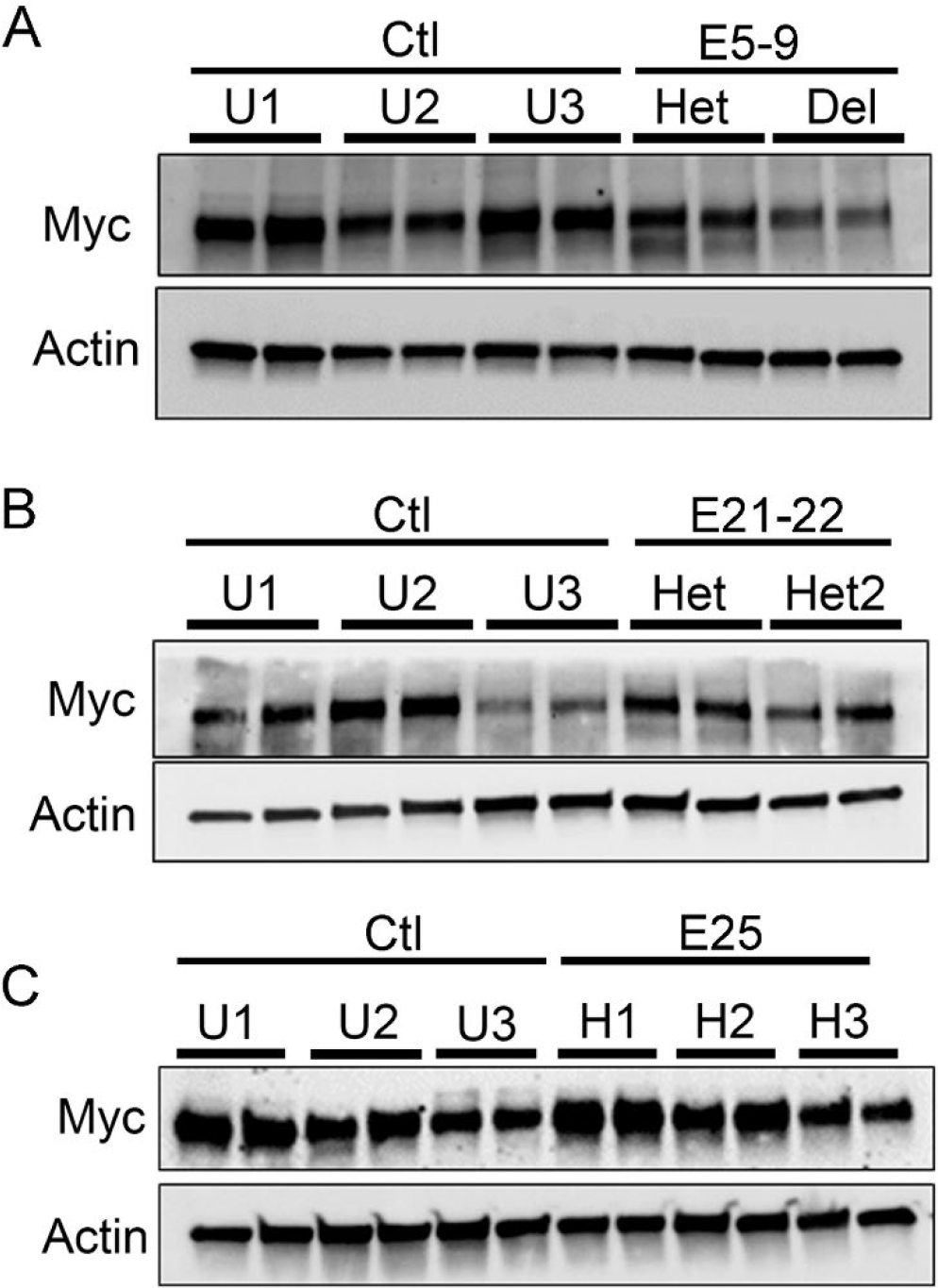
**A, B and C.** Western blot for *c-Myc* expression for loci E5-9, E21-22, and E25. P = parent control cell line. U = unedited cell line, H = heterozygous cell line Del = homozygous deletion.

**Supplementary Table 1.** Description of enhancers in the *c-Myc* locus.

| Gene Name | chrom | peak start | peak stop | peak width (bp) | Fold Change (Log2) | P value | Conserved | Previously Reported |
| --- | --- | --- | --- | --- | --- | --- | --- | --- |
| E1 | chr15 | 60982809 | 60983137 | 329 | 1.58 | 0.0494 | yes | no |
| E2 | chr15 | 61157816 | 61158677 | 862 | 0.96 | 0.0418 | yes | no |
| E3 | chr15 | 61987424 | 61988305 | 882 | 1.59 | 0.00112 | yes | yes |
| E4 | chr15 | 62039262 | 62039689 | 428 | 1.65 | 0.0303 | yes | yes |
| E5 | chr15 | 62117839 | 62118709 | 871 | 1.64 | 0.00311 | yes | yes |
| E6 | chr15 | 62118807 | 62120422 | 1616 | 2.56 | 5.07E-08 | yes | yes |
| E7 | chr15 | 62120700 | 62121878 | 1179 | 1.94 | 0.00143 | yes | yes |
| E8 | chr15 | 62122325 | 62125270 | 2946 | 2.14 | 0.00000367 | yes | yes |
| E9 | chr15 | 62127781 | 62129102 | 1322 | 2.02 | 0.0000158 | yes | yes |
| E10 | chr15 | 62129368 | 62129734 | 367 | 1.6 | 0.034 | no | no |
| E11 | chr15 | 62142375 | 62142700 | 326 | 1.63 | 0.032 | yes | yes |
| E12 | chr15 | 62157061 | 62157772 | 712 | 1.69 | 0.0162 | yes | yes |
| E13 | chr15 | 62202823 | 62204455 | 1633 | 1.44 | 0.00158 | no | no |
| E14 | chr15 | 62204551 | 62205624 | 1074 | 1.84 | 0.000596 | yes | no |
| E15 | chr15 | 62207064 | 62207770 | 707 | 1.95 | 0.00795 | no | no |
| E16 | chr15 | 62208443 | 62208807 | 365 | 2.41 | 0.00512 | no | no |
| E17 | chr15 | 62264530 | 62265748 | 1219 | 1.17 | 0.0422 | no | yes |
| E18 | chr15 | 62281950 | 62282826 | 877 | 1.28 | 0.026 | yes | no |
| E19 | chr15 | 62285294 | 62288292 | 2999 | 1.52 | 0.0000196 | yes | no |
| E20 | chr15 | 62288424 | 62289166 | 743 | 2.01 | 0.00281 | no | no |
| E21 | chr15 | 62294666 | 62295958 | 1293 | 1.03 | 0.0148 | yes | no |
| E22 | chr15 | 62298809 | 62299565 | 757 | 1.9 | 0.0121 | yes | no |
| E23 | chr15 | 62300495 | 62300964 | 470 | 2.13 | 0.00183 | no | no |
| E24 | chr15 | 62301643 | 62302876 | 1234 | 1.58 | 0.000667 | yes | no |
| E25 | chr15 | 62327111 | 62327978 | 868 | 2.15 | 0.00328 | yes | yes |
| E26 | chr15 | 62395555 | 62398548 | 2994 | 1.2 | 0.00227 | yes | no |
| E27 | chr15 | 62500301 | 62501070 | 770 | 2.92 | 0.00000627 | yes | no |
| E28 | chr15 | 62501314 | 62501898 | 585 | 2.64 | 0.00024 | yes | no |
| E29 | chr15 | 63194175 | 63194585 | 411 | 2.34 | 0.0029 | yes | yes |
| E30 | chr15 | 63484511 | 63485008 | 498 | 1.6 | 0.0279 | yes | yes |

**Supplementary Table 2.**
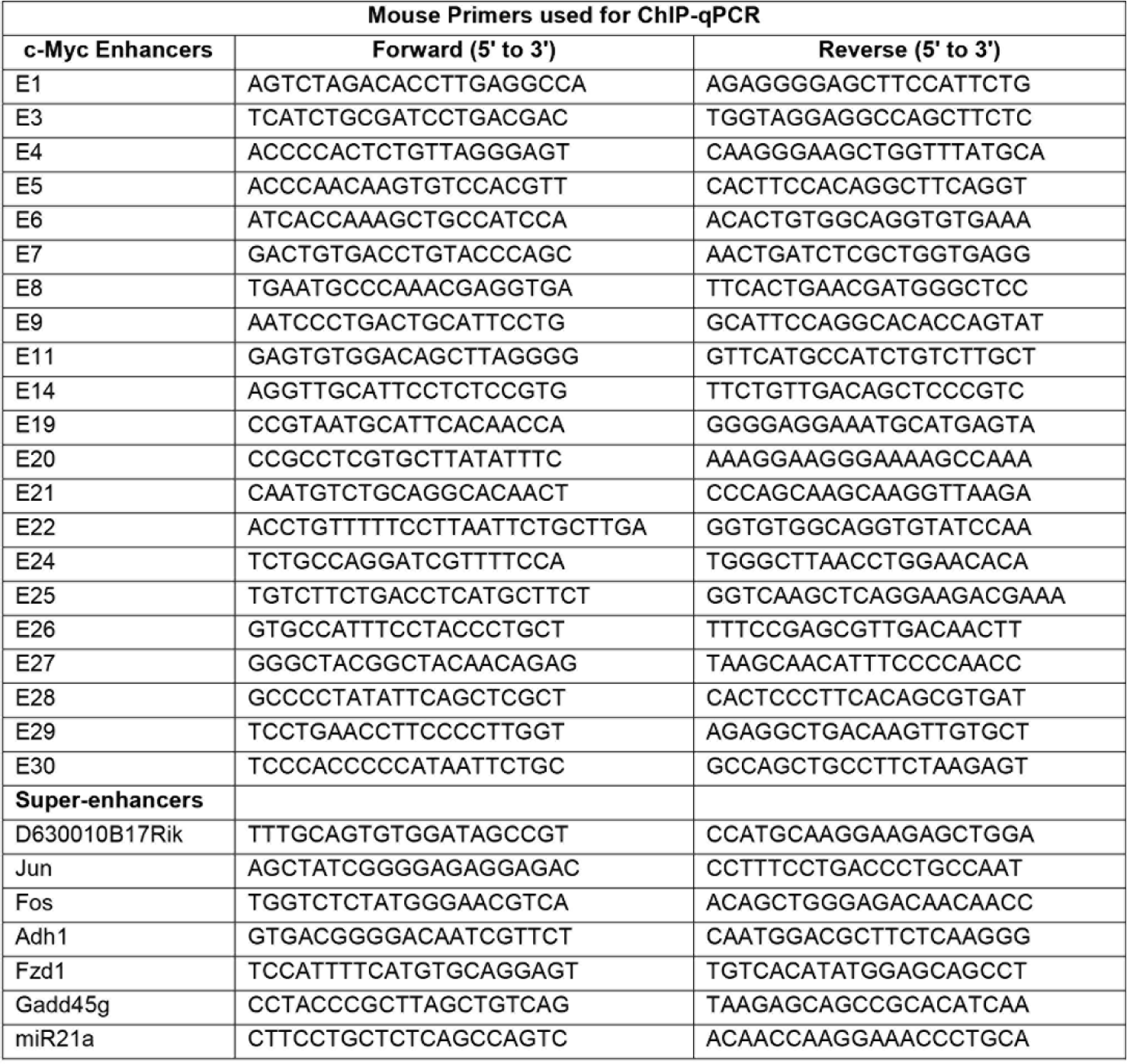
Primer sequences for ChIP-qPCR in mouse kidney samples.

**Supplementary Table 3.**
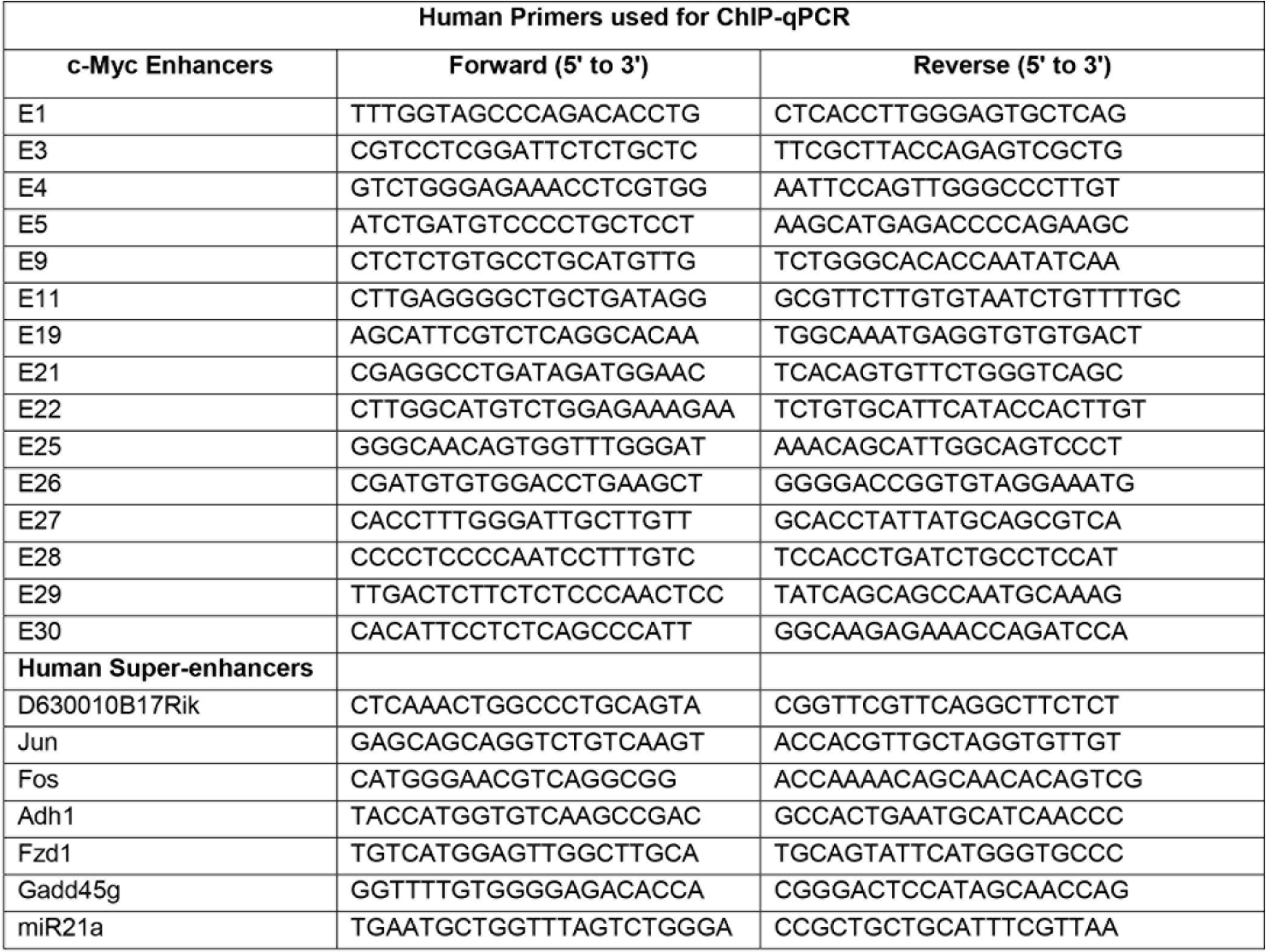
Primer sequences for ChIP-qPCR in human kidney samples.

**Supplementary Table 4.**
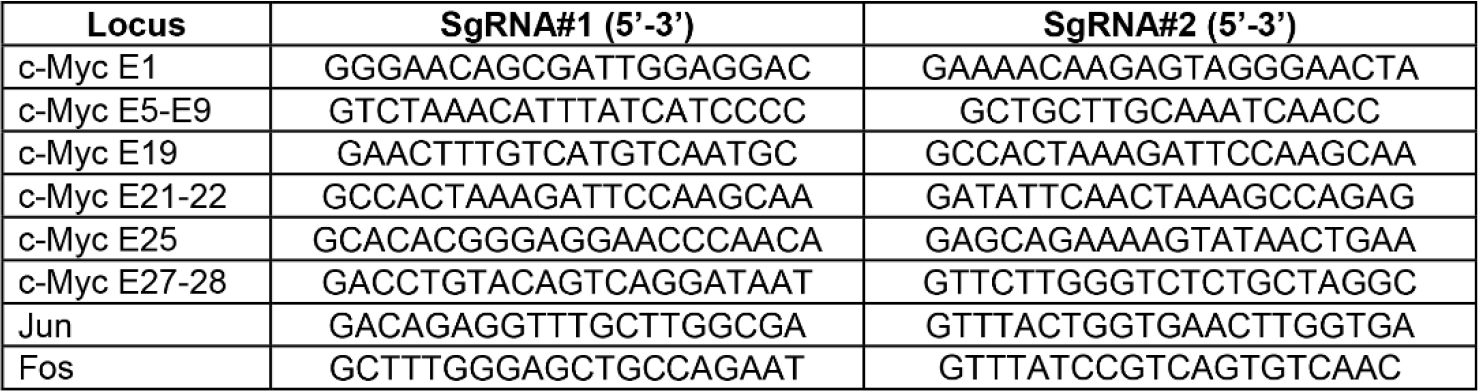
Guide RNAs used for CRISPR/Cas9 based deletion of enhancers and super-enhancers

**Supplementary Table 5.**
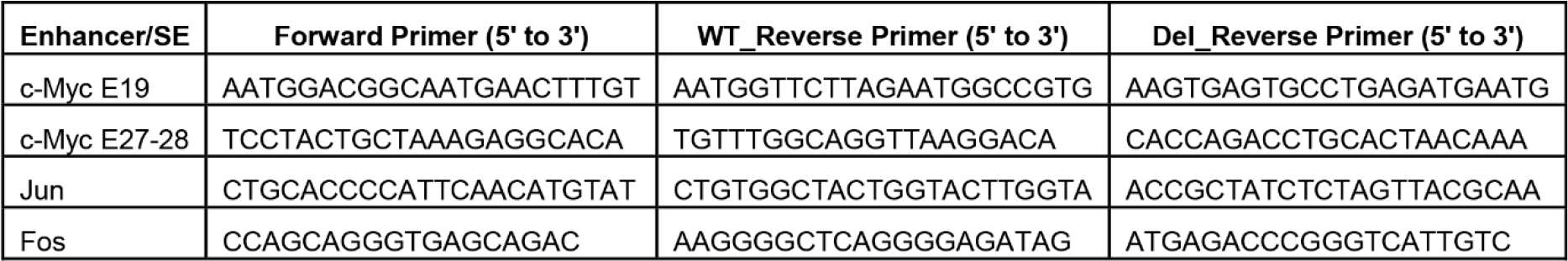
Primer sequences used to confirm deletion of enhancers and super-enhancer loci.

## Notes

### Competing Interest Statement

The authors have declared no competing interest.

